# Time between learning events shapes functional connectivity in spatial memory, a brain-wide analysis

**DOI:** 10.64898/2026.02.12.705541

**Authors:** Tommaso Gosetti di Sturmeck, Sebastiano Bergamo, Valentina Mastrorilli, Annarita Patrizi, Davide Nuzzi, Giovanni Pezzulo, Gino Del Ferraro, Arianna Rinaldi, Andrea Mele

## Abstract

A substantial body of research indicates that spaced training, characterized by longer inter-trial intervals between training epochs, consistently outperforms massed training in promoting durable memory. To investigate the neural mechanisms underlying this difference, we quantified c-Fos expression across 126 brain regions and mapped network activity following both spatial and cue-based learning under massed and spaced training protocols. While both training regimens (spaced and massed) and memory types (spatial and cue-based) produced small-world networks with similar overall topology, they differed in the functional organization of specific circuits. Massed spatial training preferentially activated hippocampal-thalamic and claustrum-basal ganglia-thalamic pathways. In contrast, spaced spatial training promoted stronger cortico-thalamic interactions and enhanced communication between the hippocampus and basal ganglia, indicating a shift toward a more integrated, cortically mediated network. These findings suggest that temporal spacing of training reorganizes memory-related brain networks, enhancing cortical-thalamic dynamics to support more efficient spatial memory. Finally, spaced networks were more sensitive to targeted disruptions of key connector hubs—identified through betweenness centrality analysis—than massed networks, pointing to a potential systems-level trade-off.

## Introduction

Training that incorporates long intervals between sessions—known as *spaced* or *distributed* training—has long been recognized as more effective for long-term retention than *massed* training, which involves short intervals between sessions^1,2^. One of the most striking aspects of this somewhat counterintuitive phenomenon—given that increasing the delay between learning and recall typically leads to greater forgetting—is its ubiquity across a wide range of learning tasks, materials, and even species^3–8^.

However, despite being first identified in the late 19th century, and despite its generalizability and potential relevance for treating individuals with intellectual disabilities, the neurobiological mechanisms underlying the spacing effect remain poorly understood. Although various theoretical models have been proposed by psychologists—each supported by experimental evidence—there is still no consensus on a single hypothesis that fully explains this phenomenon^1,5,9,10^. Among these, the model most consistent with our understanding of the biological mechanisms underlying memory formation suggests that spaced learning is effective because each subsequent retrieval adds to partially consolidated memory traces. In contrast, massed sessions do not produce the same effect, as the original memory traces remain fully active, potentially producing interference^2^. This view finds support from studies demonstrating the impact of the timing rules of learning on molecular mechanisms underlying synaptic plasticity^11–16^. Indirect support comes also from computational models that have been shown to successfully predict the efficiency of spaced stimuli in inducing plasticity in Aplysia, taking into account the complexity of the temporal dynamic of the multifaceted biochemical cascades involved^17^. Overall, this body of evidence supports the hypothesis that the superior effectiveness of spaced training over massed training in enhancing memory may be attributed, at least in part, to its impact on consolidation processes^2,12,18^.

However, efficient information processing cannot be explained solely by activity within individual brain regions; rather, it depends critically on how different regions communicate and integrate their activity. It is now well established that cognitive functions depend on the coordinated activity of widespread neural networks spanning multiple cortical and subcortical regions^19–21^. This view is supported by experimental studies using brain-mapping techniques, including analyses of metabolic changes and activity-regulated gene expression that demonstrate distributed experience-dependent changes in brain activity^22–30^. The application of graph-theoretical tools enables the characterization of functional brain networks, that can be inferred based on correlation between brain regions (nodes) activity, highlighting differences in connectivity patterns, community structure, and hub formation, thereby improving our understanding of how the brain organizes information processing to support complex cognitive functions^20,21^. Importantly, it has been found that brain functional connectivity can change depending on task demand. For example, prior learning that enhances the ability to acquire new information is accompanied by increases in brain network efficiency and architecture^31^.

Notably, a recent imaging study demonstrated differential involvement of the anterior and posterior hippocampus in the encoding of face–scene pairs following massed or spaced learning in humans^32^. Similarly, in rodents, differences in the activation of subregions within the dorsal striatum were observed during the recall of spatial information acquired in the Morris Water Maze (MWM), depending on whether massed or spaced training protocols were used^33^. Additionally, studies on memory athletes have revealed that they recruit more extensive neural circuits compared to controls, underscoring the potential role of differential network engagement in optimizing memory performance^34^. Collectively, these findings suggest that the acquisition and storage of spatial information may involve the recruitment of distinct brain circuits, depending on the characteristics of the training protocol.

To test this hypothesis, we assessed whole-brain c-Fos expression in mice immediately after training in the Morris Water Maze (MWM), using either a massed or spaced training paradigm—the latter of which has recently been shown to produce more durable memory traces^33,35^. Based on these data, we calculated inter-regional correlations of c-Fos expression and constructed functional connectivity network maps to investigate whether, and how, training frequency modulates brain networks architecture and topology during memory consolidation.

## Results

### C-fos expression after massed and spaced training

To compare regional brain activation following massed versus spaced spatial learning, we employed the Morris Water Maze (MWM) using two groups of mice trained with identical numbers of sessions but differing intertrial intervals^33^. The massed training group completed six consecutive sessions with intersession intervals of 10–15 minutes. In contrast, the spaced training group received two sessions per day over three consecutive days, with a 4-hour interval between daily sessions^33,35^. Two additional groups were trained in the cued version of the task, also over six sessions, differing only in the intertrial interval to mirror the massed and spaced training regimens used in the spatial condition (Fig. 1).

**Figure 1.**
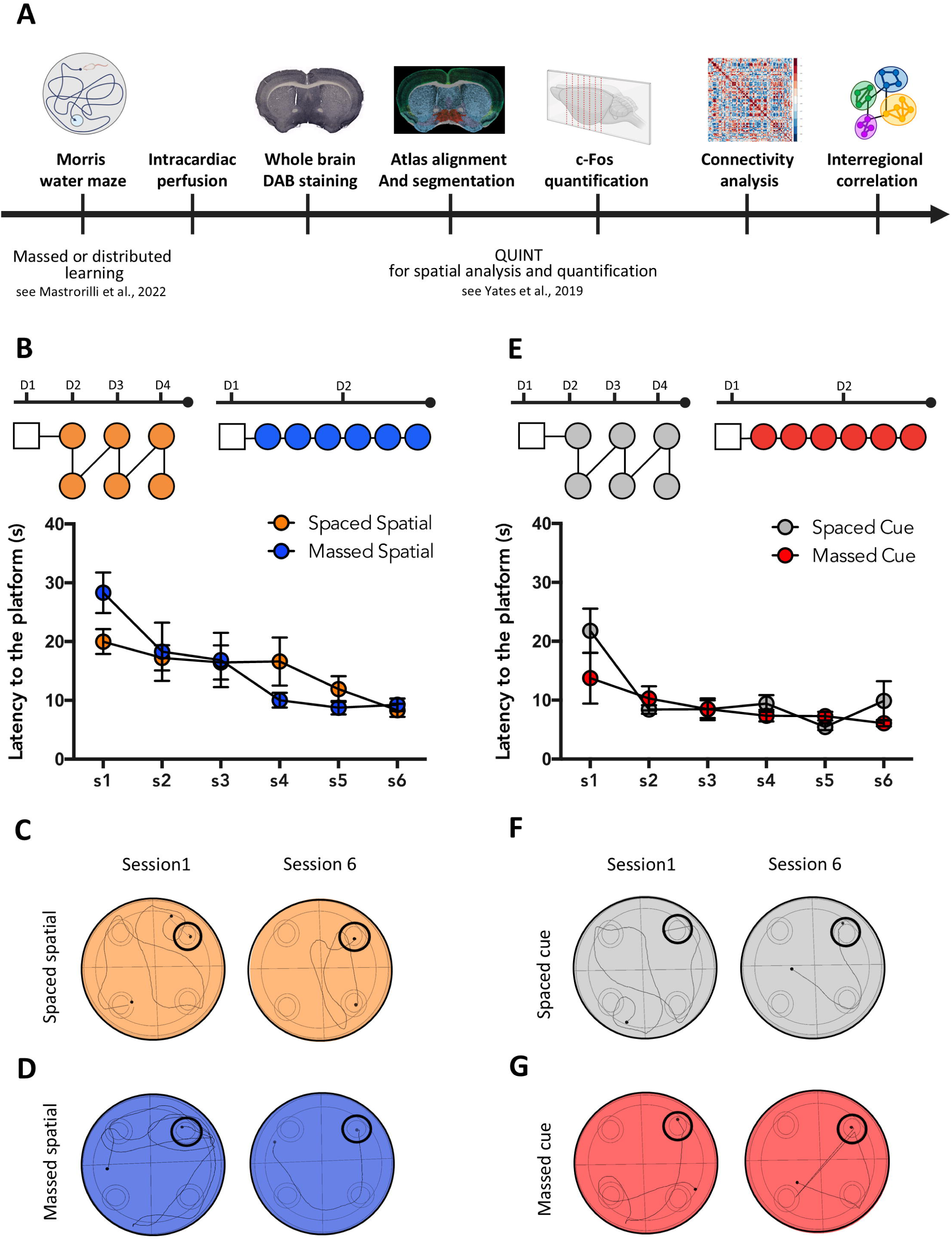
Experimental design and training protocols. **A.** Schematic of the experimental design. **B.** Schematic of the training protocol for the spatial version of the maze. **C.** Latency to find the platform for the mice trained with the spaced and massed protocol in the spatial version of the task (mean +/- S.E.M). **D.** Animals’ tracking in session 1 and session 6 after massed and spaced spatial training. **E.** Schematic of the training protocol for the cue version of the maze. **F.** Latency to find the platform for mice trained with the spaced and massed protocol in the cue version of the task (mean +/- S.E.M.). **G.** Animals’ tracking in session 1 and session 6 after massed and spaced cue training.

In the spatial MWM, mice trained under both protocols progressively reduced their latency (Fig. 1) and path length (Fig. S1) to reach the hidden platform, with no significant differences observed between groups in the learning curves or final performance levels (two-way repeated measures ANOVA for latency: session F(5,110) = 8.703, p < 0.0001; protocol F(1,22) = 0.2250, p = 0.64; session × protocol F(5,110) = 0.3482, p = 0.8825). (Fig. 1B;C;D; Fig. S1). The one-way ANOVA on time to reach the platform confirmed that both groups successfully learned the task (latency to platform: massed spatial F(5, 54) = 5.741, p=0.0003; spaced spatial F (5,72) = 4.971, p = 0.0006). Similarly, mice trained in the cued MWM showed decreased latency and distance traveled across sessions, irrespective of training protocol (two-way repeated measures ANOVA for latency: session F(5,115) = 11.76, p < 0.0001; protocol F(1,23) = 0.04857, p = 0.8275; session × protocol F(5,115) = 0.8497, p = 0.5174) (Fig. 1E;F;G). The one-way repeated measures ANOVA further confirmed that both groups acquired the task (latency to platform: massed cue F(5,66) = 3.619, p = 0.0059; spaced cue F(5, 72) = 7.937, p < 0.0001).

To assess regional neuronal activation associated with each training condition, we quantified c-Fos expression across 126 brain regions using the QUINT workflow^36,37^ in mice trained with either massed or spaced protocols in both the spatial and cue versions of the MWM. A home cage (HC) control group was also included. The analyzed regions ranged from the olfactory areas to midbrain structures, as defined by the Allen Mouse Brain Atlas^38^ (for a complete list of regions, see Table S1).

To identify regional changes in c-Fos expression under different experimental conditions, we first compared each experimental group to the home cage (HC) controls. Neural signatures of the training protocols, revealed through Partial Least Squares (PLS) analysis using a |1.96| salience threshold (p < 0.05), showed significant differences in c-Fos expression across all trained groups relative to HC controls (spaced spatial: p = 0.016; massed spatial: p = 0.00013; spaced cue: p = 0.00053; massed cue: p = 0.001) (Fig. 2).

**Figure 2.**
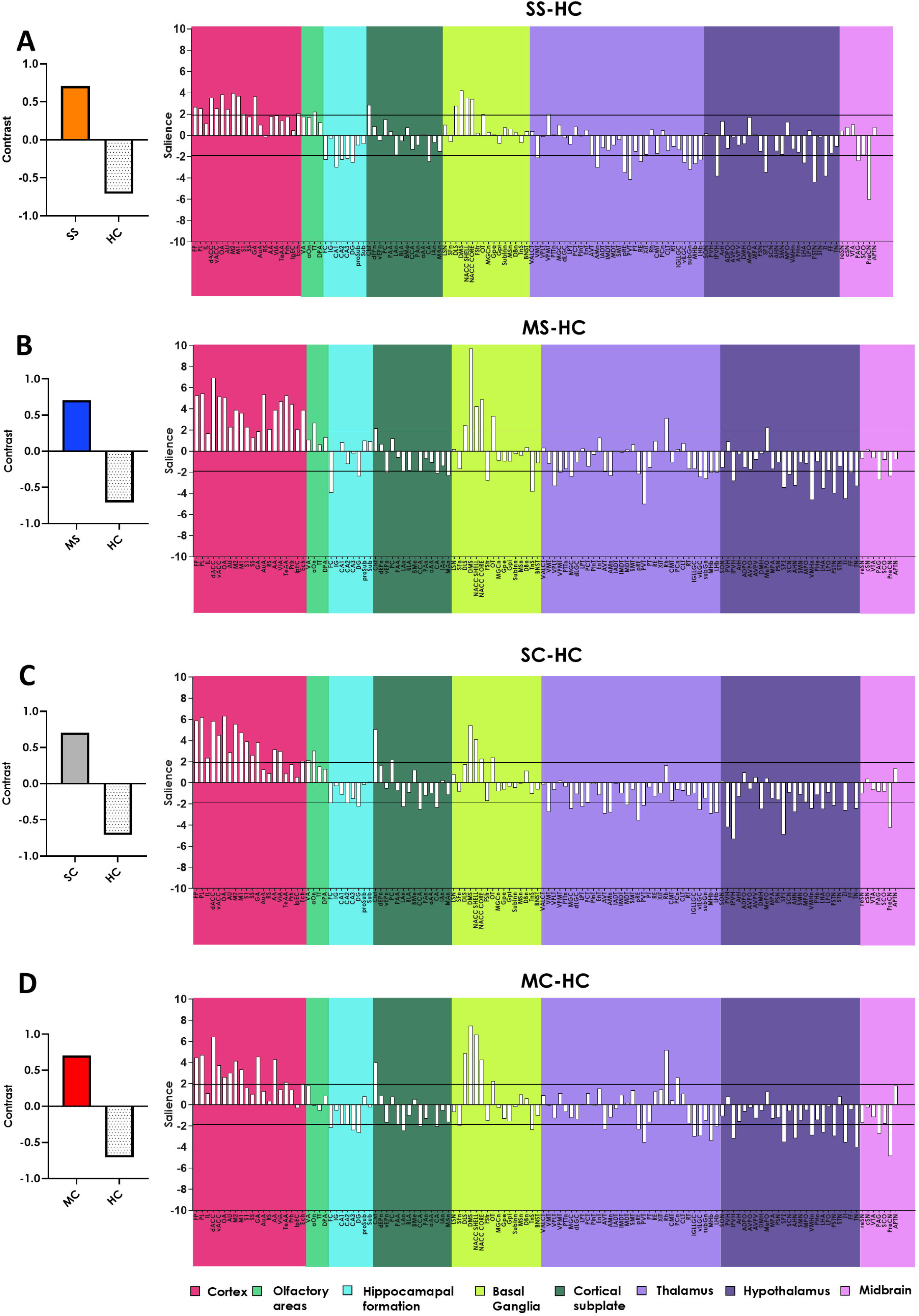
Partial least squares (PLS) analysis of c-Fos activation. PLS analysis of Fos expression across all 126 brain regions in trained vs. home cage (HC) control mice. The **left panels** show contrast analysis which robustly discriminates all trained groups from the HC controls. Spaced spatial (SS, orange), massed spatial (MS, blue), spaced cue (SC, grey) and massed cue (MC, red). The **right panels** display salience scores for individual brain regions, highlighting those that most reliably differentiate each trained group from controls (salience threshold |1.96|, p < 0.05) after spaced spatial training (**A**), massed spatial training (**B**), spaced cue training (**C**), massed cue training (**D**). Brain regions were divided into 8 major taxonomy groups: cortex (magenta), olfactory regions (light green), hippocampal formation (cyan), cortical subplate (dark green), cerebral nuclei (lime), thalamus (lavender), hypothalamus (purple), and midbrain (violet) from left to right.

The latent variables (LV) extracted by the PLS analysis captured the network of brain regions engaged by training in each group (Fig. 2), highlighting both shared and distinct patterns of activation. All trained groups exhibited increased activation in cortical and striatal regions compared to HC controls. Notably, the SS group showed a balanced engagement of motor (M1, M2) and prefrontal (PL, aCC) cortices (Fig. 2A), while in the MS group, activation was more prominent in the prefrontal cortex (Fig. 2B). Moreover, although striatal activation was generally higher in the MS group, the dorsolateral striatum (DLS) showed comparable activation in both MS and SS groups, consistent with previous findings^33^. Interestingly, the SS group also exhibited deactivation in most hippocampal subregions and in the nucleus reuniens (Fig. 2A), a pattern not observed following massed spatial training, while the rhomboid nucleus of the thalamus was highlighted only in the massed groups (Fig. 2B).

Next, we directly compared all four training conditions—spaced spatial (SS), massed spatial (MS), spaced cue (SC), and massed cue (MC)—to determine the relative contributions of training protocol and task type. PLS analysis (Fig. 3) revealed that experimental groups clustered by both task type and training protocol, with task type (spatial vs. cue) accounting for a greater proportion of the variance (LV1: 41.32%; Fig. 3A) than training protocol (LV2: 34.20%; Fig. 3B). These results suggest meaningful patterns associated with both task type and training schedule, despite not meeting conventional thresholds for significance (p = 0.236).

**Figure 3.**
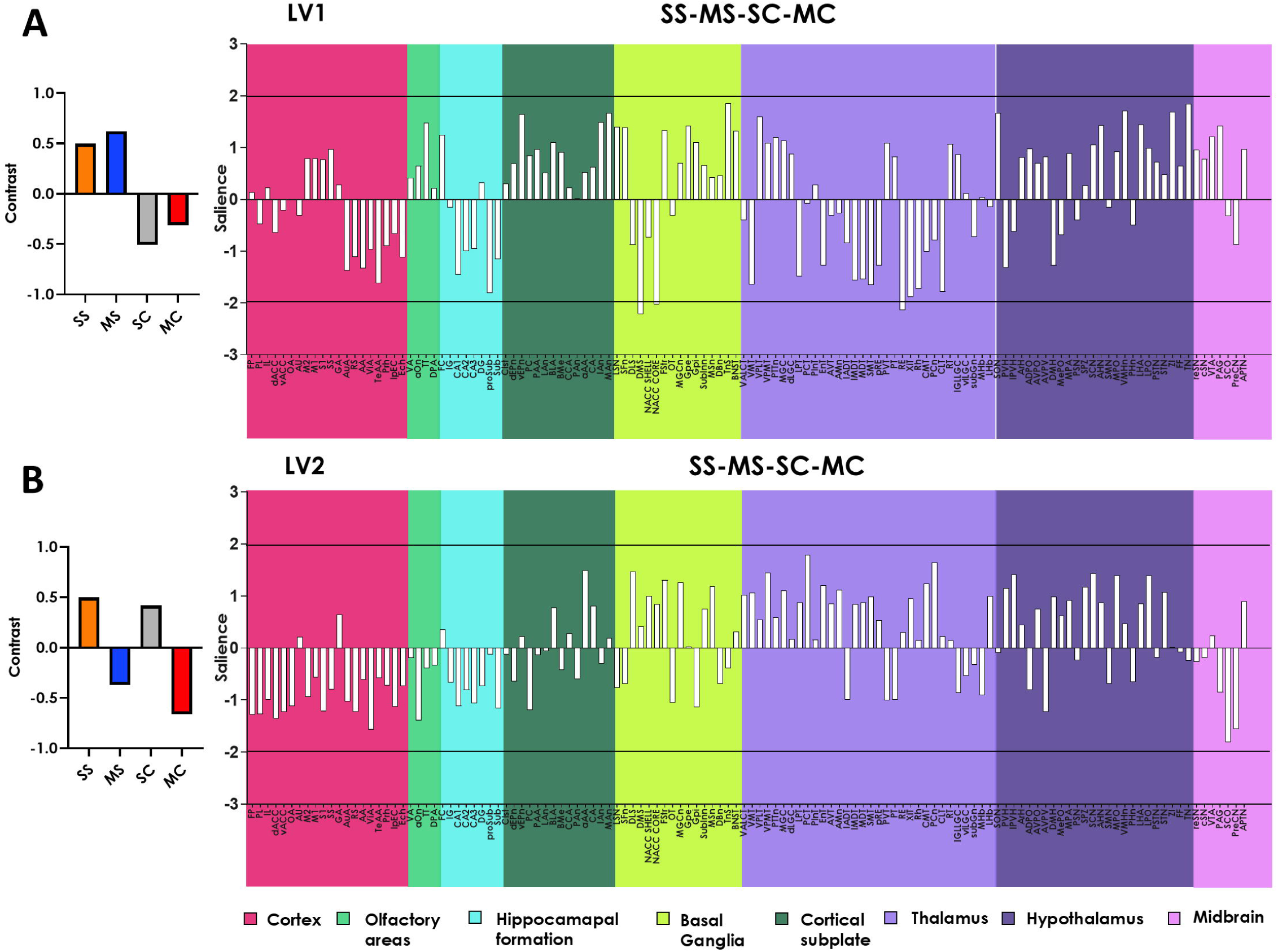
Multivariate PLS analysis distinguishes neural signatures of training protocol and task type. PLS analysis of Fos expression across all 126 brain regions in mice trained with the two different protocols (spaced and massed) in the spatial and in the cued version of the task. **A.** PLS analysis latent variable 1 (LV1) primarily separates mice based on task type, separating spatial from cued learning groups. Spaced spatial (SS, orange), massed spatial (MS, blue), spaced cue (SC, grey) and massed cue (MC, red). **B.** LV2 differentiates mice according to the training protocol, distinguishing massed from spaced training groups. In both panels, the right-side shows salience scores for individual brain regions, indicating those that reliably contribute to the group separation (salience threshold |1.96|, p < 0.05). Brain regions were divided into 8 major taxonomy groups: cortex (magenta), olfactory regions (light green), hippocampal formation (cyan), cortical subplate (dark green), cerebral nuclei (lime), thalamus (lavander), hypothalamus (purple), and midbrain (violet) from left to right.

In conclusion, our findings indicate that each training condition elicits a distinct pattern of c-Fos activation compared to home cage controls. Moreover, differences in c-Fos expression between experimental groups are primarily driven by the type of learning task (spatial vs. cue), with a smaller yet detectable influence of the training protocol (massed vs. spaced).

### Functional networks demonstrate small word topology in all experimental groups

The acquisition and storage of new information are believed to involve the coordinated activation of multiple brain regions^29,30^. While it is well established that increasing inter-trial intervals enhances memory stability, the impact of this manipulation on neural network activation patterns has been less thoroughly explored^33^. To investigate this, we examined how c-Fos activity co-varied across brain regions by computing comprehensive inter-regional correlation matrices for each experimental group, as well as for the HC controls (Fig. S2). From these correlations, we generated functional network graphs based on significant positive correlations (Pearson’s r > 0.65, p < 0.04) (Fig. 4; Fig. S3).

**Figure 4.**
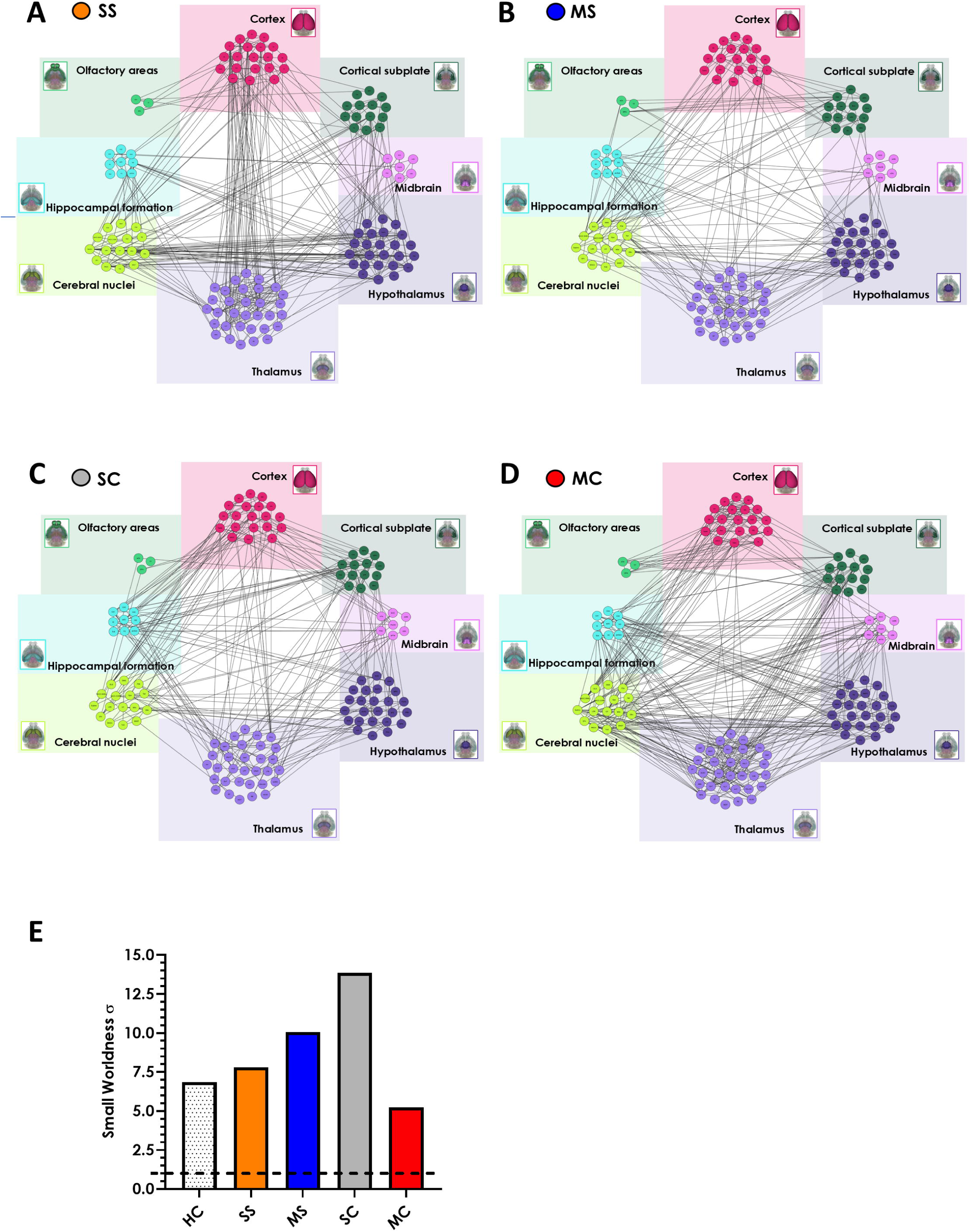
Functional connectivity networks based on c-Fos co-activation. Network graphs generated by considering only significant positive inter-regional correlations (Pearson’s r > 0.65, p < 0.04) for the spaced spatial (**A**), the massed spatial (**B**), the spaced cue (**C**) and massed cue (**D**) groups. The 126 brain regions were divided into 8 taxonomic groups: cortex (magenta), olfactory regions (light green), hippocampal formation (cyan), cortical subplate (dark green), cerebral nuclei (lime), thalamus (lavander), hypothalamus (violet), and midbrain (pink). **E.** Comparison of experimental and randomized networks demonstrates that all groups, including HC controls, exhibit small-world network properties (σ >1). Spaced Spatial (SS); Massed Spatial (MS); Spaced Cue (SC); Massed Cue (MC); Home Cage controls (HC).

We first validated the stability of these networks by comparing results using alternative correlation coefficients and assessing the reproducibility of commonly identified nodes across more and less conservative thresholds (Fig. S4A). Importantly, differences in network organization across conditions could not be attributed to artifacts from group-level differences in signal strength or variance (Fig. S4B). To assess whether standardized c-Fos density influenced the resulting correlations within our networks we compared Fos density with r^2^, a measure of correlation strength, revealing no significant correlation (p < 0.05). These analyses were conducted encompassed all 126 regions, ensuring that observed differences were not merely artifacts of group variations in signal strength and/or variance (Fig. S4B).

Although our analysis focused on positive correlations, we also observed a substantial number of negative correlations across all experimental conditions (Fig. S5), in some cases matching or exceeding the number of positive correlations, suggesting meaningful inhibitory or opposing activity patterns within the networks.

Next, we assessed whether the functional networks exhibited a small-world topology—a characteristic of biological networks that supports both functional segregation (through tightly connected local clusters) and integration (via short paths between nodes that facilitate efficient global communication)^21^. Comparisons with randomized networks revealed that all five groups displayed properties consistent with small-world organization (Fig. 4E).

We next compared the network properties of spaced-trained groups (SS and SC) to those of massed-trained groups (MS and MC). To do so, we performed a bootstrapping analysis with 1,000 iterations for each network metric. These included measures of segregation — clustering coefficient, transitivity —as well as measures of integration—global efficiency, characteristic path length (Fig. S6). This resampling approach allowed for robust statistical comparisons, accounting for network variability.

These analyses demonstrated that all networks, regardless of training protocol or task type, shared a comparable small-world topology (Fig. 4E) and similar values for key integration and segregation metrics (Fig. S6). Since no significant topological differences emerged at this global level, we proceeded to a more detailed analysis of individual network features, with the goal of identifying specific patterns of connectivity that might underlie the distinct effects of massed versus spaced training.

### Spatial memory networks depend on the training protocol

To test this hypothesis, first we performed a brain-wide comparison of inter-regional correlations, grouping brain areas into 20 anatomical and functional clusters to create coarse-grained networks (Table S1). With few exceptions (see methods) clusters were delineated accordingly to the Allen Brain Atlas^38^. For further improved clarity regarding networks interpretation, we have chosen to exclude connectivity arising from connections with the hypothalamic regions at this stage. By reducing the dimensional complexity of interregional interactions, this approach revealed significant differences in functional connectivity patterns between networks generated by massed and spaced spatial training (Fig. 5).

**Figure 5.**
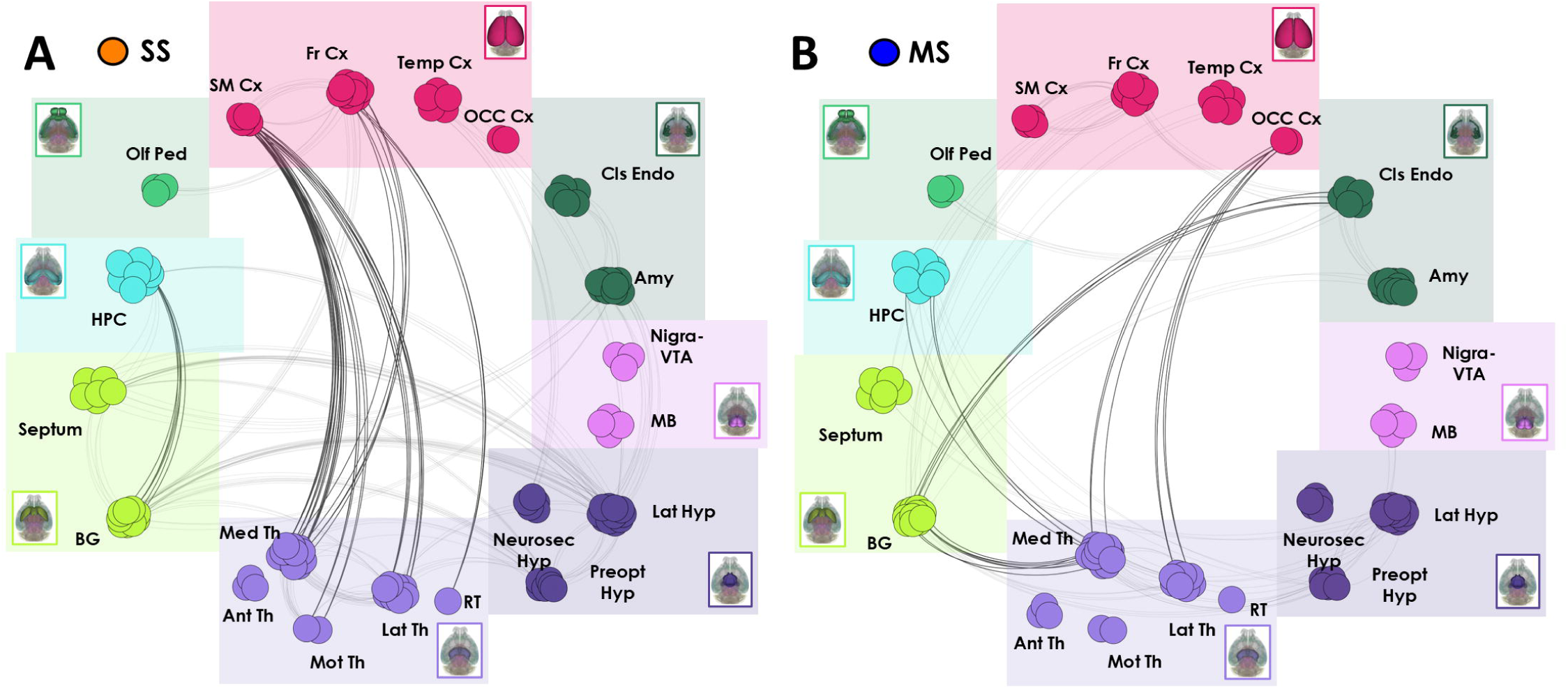
Coarse grained functional connectivity networks in spaced spatial (SS) and massed spatial (MS) training groups. Brain networks, based on c-Fos correlation patterns. The 126 brain regions divided into 8 taxonomic groups, were organized into 20 functional-clusters (cortex (magenta): somato-motor cortex (SM Cx), frontal cortex (Fr Cx), temporal cortex (Temp Cx) and occipital cortex (OCC Cx). Olfactory regions (light green): olfactory peduncle (Olf Ped). Hippocampal formation (cyan): hippocampus (HPC). Cerebral nuclei (lime): Septum and basal ganglia (BG). Thalamus (lavender): medial thalamus (Med Th), anterior thalamus (Ant Th), motor thalamus (Mot Th), lateral thalamus (Lat Th), reticular thalamus (RT). Hypothalamus (purple): neurosecretory hypothalamus (Neurosec Hyp), preoptic hypothalamus (Preopt Hyp), and lateral hypothalamus (Lat Hyp). Midbrain (violet): nigra and ventral tegmental area (Nigra-VTA) and midbrain regions (MB). Cortical subplate (dark green): claustrum and endopiriform regions (Cls Endo) and amygdala (Amy). Edges are represented as connections between clusters only if there were at least three significant inter-cluster connections, highlighting robust communication. Darker lines denote interesting relevant correlations. **A.** SS exhibited stronger correlations between the frontal cortex and medial thalamus and between the hippocampus and basal ganglia. **B.** Conversely, strong functional connectivity between both the hippocampus and the basal ganglia with the medial thalamus was consistently observed in MS. Spaced Spatial (SS); Massed Spatial (MS)

In particular, the network associated with massed training (MS) showed strong connectivity between allocortical and subcortical structures, particularly with the thalamus. In contrast, the spaced training (SS) group is characterized by lower connectivity between subcortical regions, while cortico-thalamic connectivity becomes more prominent (Fig. 5A;B). Specifically, the MS network exhibited robust interregional correlations between the hippocampal formation and the medial thalamic nuclei, as well as between the basal ganglia and the medial thalamus (Fig. S7). Conversely, the SS network displayed stronger functional connectivity between the somatosensory cortex and medial, motor, and lateral thalamic nuclei, along with notable connections between the frontal cortex, the medial thalamus, and the thalamic reticular nucleus (Fig. 5A; Fig. S8). Additionally, in the MS network, we observed strong correlations between the claustrum and the basal ganglia, and between the occipital cortex and medial thalamic nuclei—patterns not present in the SS network (Fig. S7). On the other hand, the SS network was characterized by enhanced functional connectivity between the hippocampus and basal ganglia compared to the MS network (Fig. 5A; Fig. S8).

Further comparisons across networks generated from cue-based learning using both spaced and massed protocols highlighted connectivity patterns linked to the duration of inter-trial intervals (Fig. S9). Notably, strong functional connectivity between the basal ganglia and the medial thalamus was consistently observed in massed training conditions but was absent in spaced training (Fig. S10). Conversely, spaced training networks exhibited stronger correlations between the frontal cortex and medial thalamus compared to massed protocols (Fig. S10).

Interestingly, cue learning generated unique connectivity patterns not seen in spatial learning networks. These included functional correlations between the somatosensory cortex and the basal ganglia, as well as between amygdalar nuclei and both medial and lateral thalamic regions (Fig. S11).

Together, these findings support our initial hypothesis that the neural networks activated by spatial learning are strongly influenced by the training protocol, regardless of the type of information being acquired. Furthermore, they demonstrate that increasing the interval between training trials shifts the dominant functional connectivity from hippocampal–thalamic circuits, seen in massed training, toward cortico–thalamic circuits in spaced training.

### Identifying memory essential hubs

Overall, our findings demonstrate that the acquisition of spatial information through either massed or distributed training engages broad—but distinct—neural networks, each characterized by elevated activity and unique patterns of functional connectivity. Importantly, a node’s influence within a network is not determined solely by the number of its connections (degree), but also by the nature and strategic positioning of those connections. Effective network function relies on both highly connected hub regions and bridging nodes that link distinct modules, thereby facilitating cross-network communication^21,23,39^.

To gain a more comprehensive understanding of node importance, we calculated and ranked nodes using additional graph theory metrics: Katz centrality, betweenness centrality, and Collective Influence (CI). To assess network resilience and robustness, we simulated two types of perturbations: random node removal (45 iterations) and targeted attacks based on ranked metrics. In the targeted attacks, nodes were removed sequentially according to their highest values for degree, Katz centrality, betweenness centrality, or CI (30 iterations each). After each removal, metrics were computed again on the resulting network recalculating the size of the largest connected component (i.e., collection of connected nodes) before proceeding to remove the next highest-ranked node according to the selected metric.

Our results revealed that all networks showed notable resilience to disruptions based on degree and Katz centrality, retaining more than 50% of their original giant component (GC) - representing the largest interconnected subnetwork through which information can propagate efficiently - even after the removal of at least 25 nodes (Fig. 6). In contrast, networks were more vulnerable to disruptions based on CI and betweenness centrality—especially in the distributed training networks. In spaced networks, GC size dropped by 50% after the removal of just four high-betweenness nodes, whereas in massed networks, this threshold was not reached until the removal of eight such nodes (Fig. 6). These findings suggest that while hub nodes are important across all networks, bridging nodes are particularly crucial for maintaining integration in networks formed through spaced training.

**Figure 6.**
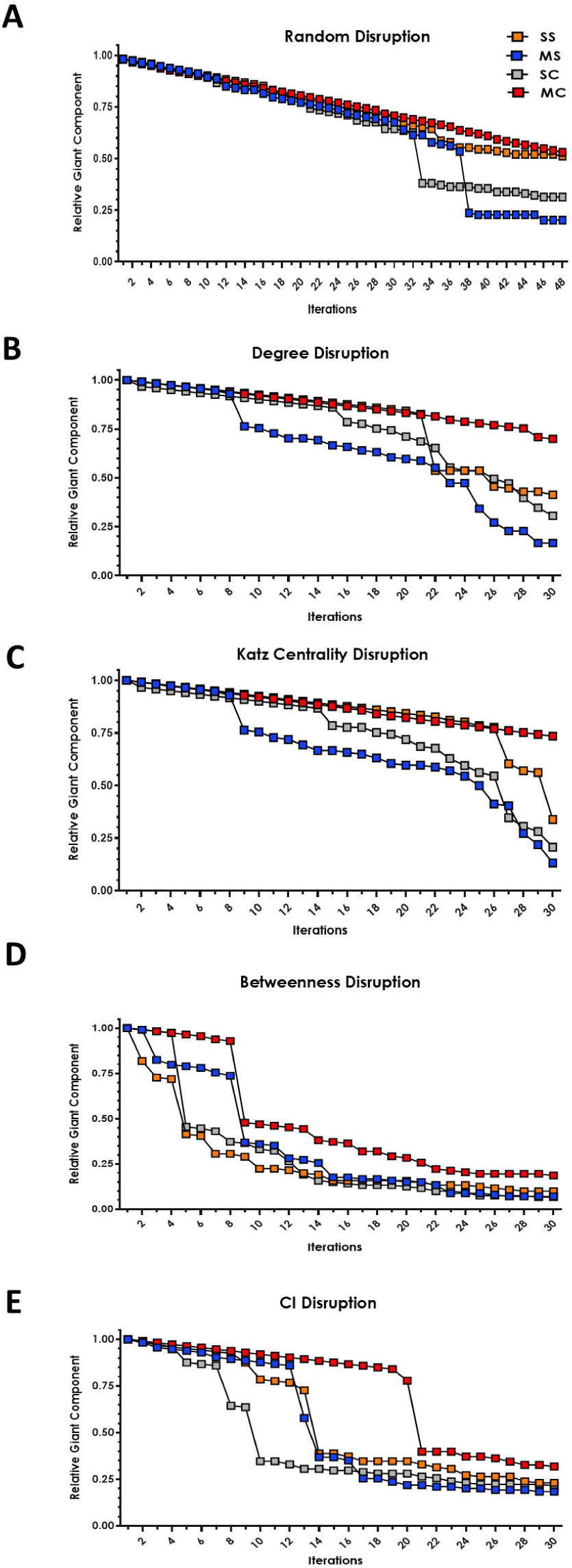
Network resilience to node removal: random deletions and targeted attacks based on centrality metrics. **A.** Effect of random node removal on the size of the largest connected component in all networks: spaced spatial (SS, orange squares), massed spatial (MS, blue squares), spaced cue (SC, grey squares), and massed cue (MC, red squares). Results were obtained via 45 bootstrap iterations, showing a progressive decline in largest component size as nodes were randomly removed. **B.–E.** Effect of targeted node removal by sequentially deleting nodes ranked in descending order according to different centrality measures: **B.** degree, **C.** Katz centrality, **D.** betweenness centrality, and **E**. Collective Influence (CI). Each targeted attack involved 30 iterations with network reconstruction after each node removal. In all panels, the size of the largest connected component is expressed as a proportion of the original largest component size.

We further organized nodes based on their rankings across the different metrics and evaluated their functional relevance using disruption-induced changes. This approach identified both highly connected hubs (via degree and Katz centrality) and bridging nodes (via CI and betweenness centrality). Although degree and Katz centrality are less precise measures of integration, they are widely used for identifying hubs^40,41^. In the spaced spatial (SS) network, hubs identified by degree were primarily located in the sensorimotor cortex and thalamus, with the paracentral nucleus of the medial thalamus ranking among the highest. In contrast, high degree nodes in the massed spatial (MS) network included distinct components of the striatal complex—such as the dorsolateral striatum, nucleus accumbens (shell), the rhomboid nucleus of the medial thalamus, and the claustrum (Fig. S12).

To identify critical nodes beyond those defined by high degree, we used betweenness centrality to estimate the minimal sets of nodes required to disrupt the GC. This metric captures a node’s influence over shortest path routing within the network^41^. These analyses confirmed several previously identified hubs but also uncovered additional critical nodes not identified by degree alone (Fig. 7). In the SS network, thalamic nuclei again emerged as central, with the paratenial and ventromedial nuclei ranking among the most critical (Fig. 7). In the MS network, the claustrum emerged as both a hub and a connector node essential for inter-cluster communication (Fig. 7). Furthermore, betweenness centrality analysis revealed key contributions of hippocampal regions—CA1 in the SS network and CA3 in the MS network—across both conditions. Unexpectedly, the SS network analysis also identified the ventral tegmental area (VTA) and substantia nigra (SN) as critical nodes, regions not observed in the MS or cue-based networks (Fig. 7).

**Figure 7.**
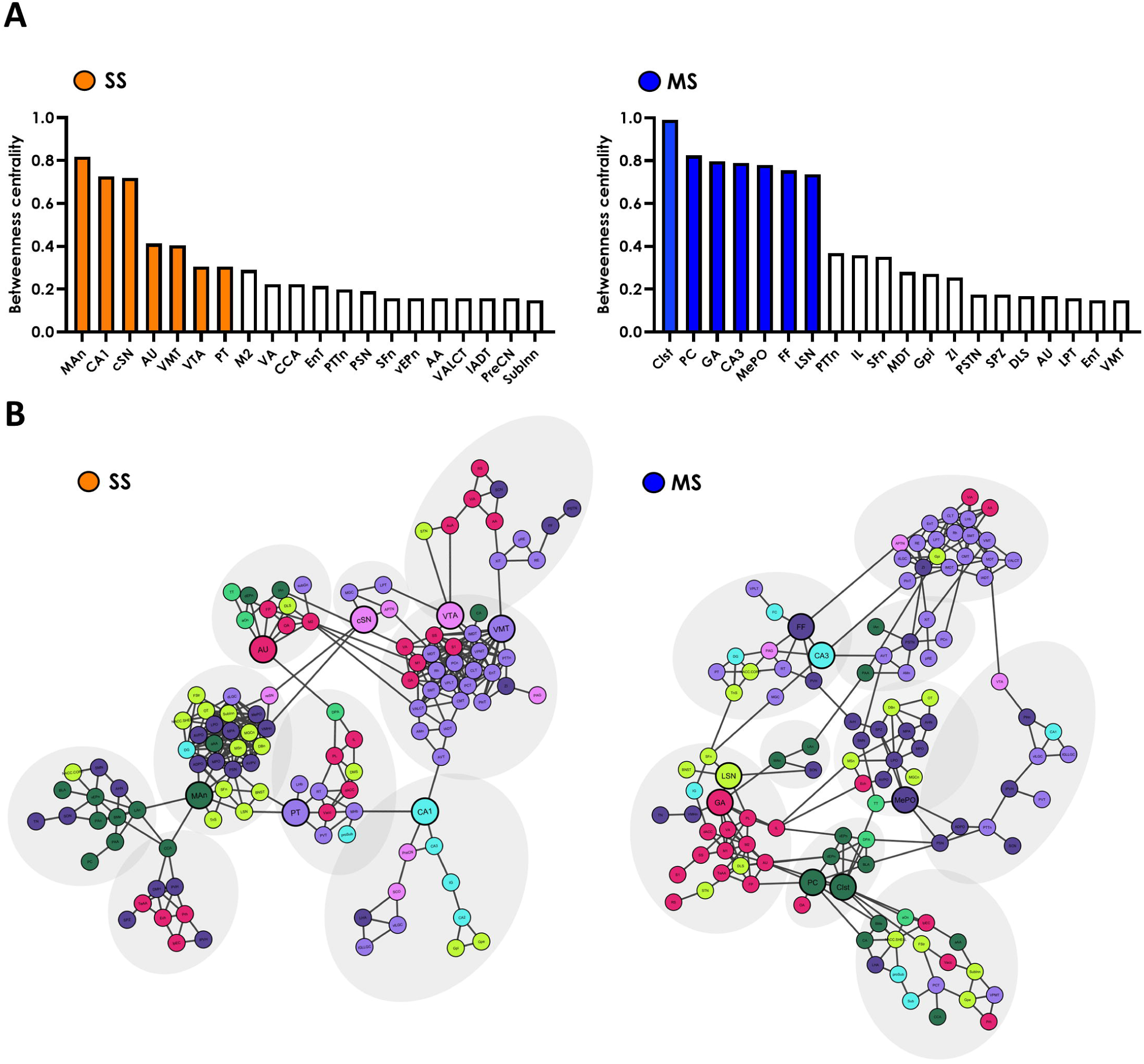
Ranking and network visualization of the most disruptive nodes in spaced and massed spatial networks based on betweenness centrality. **A.** Disruption ranking of nodes in the spaced spatial (SS, orange) and massed spatial (MS, blue) networks derived from simulated node removal using betweenness centrality. The top 7 nodes for each group are highlighted in their respective colors; all other nodes are shown in white. **B**. Network layout visualizations generated with Cytoscape for SS and MS groups. Grey ellipses highlight networks’ communities. Nodes are color-coded according to their original taxonomical group: cortex (magenta), olfactory regions (light green), hippocampal formation (cyan), cortical subplate (dark green), cerebral nuclei (lime), thalamus (lavander), hypothalamus (purple), and midbrain (violet). The top 7 ranked disruptive nodes (from panel A) emphasized by larger node sizes to highlight their network importance.

Cue learning networks exhibited distinct connectivity profiles relative to spatial networks. In the spaced cue (SC) network, the globus pallidus and lateral thalamic nuclei (medial and lateral geniculate) ranked highly, acting as integrators across modules. Notably, CA1 also ranked highly in the SC network, suggesting shared features among spaced networks regardless of learning type. In the massed cue (MC) network, the prelimbic cortex (PL) ranked prominently (Fig. S13).

Together, the identification of both hubs and bridging nodes across these networks supports conclusions drawn from broader network analyses, particularly emphasizing the centrality of thalamic nuclei—especially the paratenial and ventromedial nuclei—in the SS network. Additionally, we highlighted the important role of CA1 and dopamine-releasing nuclei in spaced spatial learning, and the involvement of basal ganglia nuclei as both hubs and connectors in massed spatial training.

## Discussion

In the present study, we employed global mapping and graph-theoretical approaches to compare massed and spaced spatial training in the Morris water maze, aiming to highlight network-level features that could explain the enhanced memory efficiency induced by longer inter-trial intervals. Furthermore, we compared massed and spaced training using both the spatial and cue versions of the task to identify commonalities and differences related to the training protocol or type of memory.

This analysis yielded several noteworthy observations. First, we found that training-induced neuronal activity in all groups engaged a wide network of brain regions, primarily involving cortical and basal ganglia areas, but with significant qualitative and quantitative differences depending on the protocol or memory type. Second, all functional networks identified by our analysis exhibited small-world topology, which is typical of biological systems. Interestingly, spaced training networks—regardless of the type of memory—were characterized by higher sensitivity to betweenness disruption, indicating greater efficiency compared to massed training networks. Finally, correlation analyses revealed substantial differences in the networks activated by spaced versus massed spatial training, suggesting a stronger engagement of cortico-thalamic loops following distributed training, compared to cortical sensory-basal ganglia correlations as well as basal ganglia-thalamic correlations, which were more prominent after massed training.

To identify network-level features associated with massed and distributed training, we generated whole-brain maps of c-Fos activity immediately after the final training session in mice trained using the two protocols. Our working hypothesis was that the neural networks engaged by distributed (spaced) training—which has been shown to strengthen memory and extend its persistence^8,33^—would differ from those recruited by massed training. Consistent with our hypothesis we found several differences between the groups trained with the two protocols. The first observation was that overall c-Fos density was higher in the massed groups compared to spaced, independently on whether mice were trained to find a visible or a hidden platform. Interestingly, the PLS analysis comparing trained groups with HC controls revealed a similar pattern: the contribution (saliency) of individual brain regions to the latent variables (LVs) was greater in the massed spatial (MS) groups than in the spaced spatial (SS) groups. These findings suggest that massed training elicits stronger region-to-region interactions or more pronounced network-level c-Fos responses than spaced training. Overall, these results parallel with previous observations demonstrating a progressive reduction in the number of c-Fos-expressing cells across training sessions^42^, which has been interpreted as a marker of increased network efficiency—wherein lower neuronal activation is required to achieve improved performance^31,42^. Interestingly the PLS-derived patterns emphasize that both training schedules engage overlapping prefrontal–striatal systems, but with differential coordination. More importantly, when PLS analysis was conducted across all groups together, LV1 and LV2 captured, respectively, the differences associated with the type of memory and the protocol used. Although these LVs did not reach statistical significance, they explained a substantial proportion of the observed variability (41% and 32%, respectively). The importance of task demands as a primary determinant of brain activation patterns is unsurprising. However, the finding that the temporal structure of training (i.e., the spacing between sessions) also shapes large-scale neural activity supports the hypothesis that massed and spaced training engage distinct neural circuits.

The functional networks derived from spaced and massed spatial training further highlighted both topological and organizational differences. Specifically, our analysis revealed that the spaced spatial network is characterized by a higher number of edges (i.e., correlations between nodes/brain regions) (Fig. 4), higher number of nodes and greater sensitivity to betweenness and CI disruption, but although to a lesser extent lower sensitivity to degree and Katz centrality disruption compared to the massed spatial network (Fig. 6 and 7). Betweenness centrality reflects the influence of nodes based on the shortest paths within the network and highlights low-degree nodes that act as bridges between larger communities^2,43,44^. Interestingly, the network generated by spaced training to locate a visible platform also shows high sensitivity to betweenness disruption (Fig. 6). Overall, these findings suggest that, compared with the massed spatial network, the spaced network is more distributed yet relies on a limited set of key nodes critical for efficient information flow. This organization suggests that the greater stability of memories formed through spaced training, could depend upon a more distributed memory trace that renders the representation more resilient to disruption.

Further and more interesting analysis of brain functional network revealed also a completely different signature in the two conditions immediately after training. Indeed, the spaced spatial network was characterized by strong cortico-thalamic correlation, with particular emphasis on the connectivity between the frontal cortex and the medial and lateral thalamus (Fig. 5A). On the contrary, the massed spatial training network was characterized by a different profile highlighting again a central role of the medial thalamus but with privileged correlation with the basal ganglia, the hippocampal complex and the occipital cortex (Fig. 5B). Interestingly, the cortical dominance observed following spaced training is consistent with the hypothesis that longer inter-trial intervals could promote incremental adjustments of a consolidated memory trace rather than interfering with representations that remain fully active, thereby supporting long-lasting storage^2^. Further, the analysis of the two networks identified different topological control hubs: a mesencephalic dopaminergic nuclei-thalamic-hippocampal axis for the spaced network while the massed network was dominated by claustral and CA3-centered control (Fig. 7A;B).

In the present study, we compared massed and spaced spatial training using the same apparatus and an identical number of sessions, with the two protocols differing only in the time intervals between training sessions. Under these conditions, previous work has shown that mice do not differ in their ability to recall the position of the platform 24 hours after the last training session^33^. Therefore, since the information to be acquired are identical in both conditions, one might expect the massed and spaced networks to display similar profiles, consistent with the prevailing view that spatial memory relies on a core set of brain regions^45–47^. Indeed, many of the regions identified in our analysis—including the hippocampus, the thalamus, the frontal cortex, the claustrum, the striatum and dopaminergic nuclei^48–54^—have been previously implicated in this core spatial memory network, providing validation for our approach. It is worth noting that the analysis revealed a greater engagement of the dorsolateral striatum (DLS) relative to the dorsomedial striatum (DMS) following spaced, compared to massed, training, consistent with previous findings^33^.

The results of our functional analyses suggest that the timing of training is a crucial determinant in shaping the consolidation network. Although differences in retrieval performance following massed versus spaced training have already been reported in both humans^32^ and rodents^33^, our findings provide the first evidence of large-scale network-level differences associated with these training schedules. It is worth noting that, although the information to be acquired is identical in both conditions, differences in task demands—such as attentional effort or the relevance of sensorimotor information—may account for some of the observed variations in network architecture. The prominent centrality of the claustrum in the massed training network, a region proposed to act as a hub for integrating sensory and limbic information to influence attention^50,55^, is consistent with this interpretation. The question remains whether and how the specific signature of the spaced network could explain the optimization of memory induced by spaced training. Based on the functional architecture of the spaced network it could be suggested that longer inter-trial-intervals allow a more efficient engagement of thalamo-cortical interplay, not observed in the MS group, that could underlie a better thalamic coordination of cortical activity, necessary for the constitution of stable, and more enduring memories induced by spaced training. The centrality of dopamine neuromodulatory regions in this network aligns with the ability of catecholaminergic inputs to shape such thalamo-cortical dynamic^56^. In contrast, the accelerated learning in the MS group promotes a basal ganglia-thalamic interaction, less efficient at stabilizing enduring memory traces.

Using a graph-theoretical approach applied to whole-brain c-Fos mapping, we demonstrate a distributed pattern of activity immediately after spatial training in the MWM, consistent with previous findings from other memory tasks^25,26,28–30^. Moreover, in agreement with prior correlational and causal evidence from both humans and rodents^32,33^, our comparison of spaced and massed training protocols revealed that the network architecture induced by learning is strongly dependent on the temporal structure of training. Finally, the topological features and brain regions identified by our analysis suggest that the enhanced memory stability observed with increased spacing between training sessions may rely on shift toward a more distributed cortically mediated network, characterized by a cortico-thalamic signature under the influence of dopaminergic hubs, whereas massed training relying on sensory motor architecture, maintains a locally efficient but less integrated configuration.

## Resource availability

### Lead contact

Requests for further information and resources should be directed to, and will be fulfilled by, the lead contact, Andrea Mele.

### Materials availability

This study did not generate new, unique reagents.

### Data and code availability

The analysis code and repository files associated with this study are available in the GitHub release v1.0.1 at https://github.com/tommasogosetti/c-Fos_mapping_massed_vs_spaced/releases/tag/v1.0.1. An archived version of the repository is available on Zenodo at https://doi.org/10.5281/zenodo.20122381.

Any additional information required to reanalyze the data reported in this paper is available from the lead contact upon request.

## Acknowledgments

We thank Matthieu Wolff, Cornelius T. Gross, and Giulia Torromino for useful discussions and suggestions, and Federico Silvestri for assistance with the analyses. We would also like to thank Chiara Feroleto for her assistance with the QUINT workflow. We thank the Light Microscopy Facility (DKFZ) for microscope use. This study was supported by a PRIN2022 Grant from MUR (to A.M.), by grants from Sapienza University of Rome (to A.M.; A.R.).

## Authors contributions

Conceptualization V.M., T.G.d.S., A.M., A.R.; behavioral experiments T.G.d.S., and V.M.; histology and microscopy A.P., V.M., T.G.d.S.; formal analysis, T.G.d.S., S.B., D.N., G.d.F., G.P.; writing – original draft, T.G.d.S., A.M.; A.P. writing – review and editing, , T.G.d.S., S.B., D.N., G.d.F., G.P., A.P. A.R., A.M, V.M..

## 3. Methods

### EXPERIMENTAL MODEL AND STUDY PARTICIPANT DETAILS

#### 3.1 Animals

Experiments were conducted on naïve CD1 male mice (Charles River). Mice were at least 11-18 weeks old, weighing about 40-60 g at the onset of the experimental procedures. Animals were housed in groups of three to five in standard cages (26.8 × 21.5 × 14.1 cm), with water and food ad libitum, under a 12-h light/dark cycle and constant temperature (22 ± 1 °C). Behavioral training was conducted during the light cycle (from 9:00 AM to 5:00 PM). All animals were treated according to current Italian and European laws for animal care, and the maximum effort was made to minimize animal suffering. Procedures were conducted under the authorization n° 658/2019 from the Italian Ministry of Health, according to Italian (DL. 26/2014) and European laws and regulations on the use of animals in research, and NIH guidelines on animal care.

### METHOD DETAILS

#### 3.2 Behavioral Procedures

Mice were trained alternatively in one of two Morris Water Maze (MWM) setups: the spatial and the cue version, both consisting of a familiarization and a training phase. Familiarization was the same for both massed and distributed protocols. Massed training consisted in six consecutive sessions (intersession interval (ISI): 10 to 15 min) of three trials (intertrial interval (ITI): 30 s), while the distributed training consisted of six sessions of three trials (ITI: 30 s) spaced over 3 days, two sessions per day (ISI: 4 h). In the spaced version, throughout training, 4 different bi-dimensional or tri-dimensional visual cues were placed on the curtains in fixed positions while in the cMWM all the distal cues were completely removed, and a proximal cue, constituted by a green ball, was hung 5 cm above the hidden platform. In the cue version the position of the platform and the ball changed across sessions to prevent animals from using spatial bias. Behavioral data from training trials were acquired and analyzed using an automated tracking system (ANY-maze, Stoelting).

#### 3.3 Immunohistochemistry: c-Fos Staining

One hour after completing the last training session, each animal was deeply anesthetized with a mixture of Zoletil 100 (100 mg/kg, Virbac Italia) and Xylazine (20 mg/kg, Bayer) and transcardially perfused with 40 ml of saline solution (NaCl 0.9%) followed by 40 ml of 4% formaldehyde. Brains were rapidly dissected and post-fixed for 24 h in 4% formaldehyde (Sigma-Aldrich, USA) in PBS and then transferred to a 30% sucrose solution in PBS (Sigma-Aldrich). 30μm-coronal sections were obtained using a freezing microtome (Leica CM 1950, Leica Microsystems, Wetzlar, Germany). Sections distanced 90 μm (one in every three) were selected to be processed for c-Fos-immunohistochemistry. Sections were transferred to 0.1 M phosphate-buffered saline (PBS, pH 7.4) and washed several times. Sections were treated for 5 min with 3% hydrogen peroxide in PBS. After 1 h of incubation with PBST containing 1% BSA and 1% NGS 3% (Normal Goat Serum) (PBST–BSA–NGS), sections were incubated overnight with anti-phospho-c-Fos rabbit monoclonal antibody (5348S; Cell signaling Technology, USA) diluted 1:8000 in PBST–BSA–NGS. The next day sections were incubated with biotinylated secondary antibody diluted 1:500 in PBST-BSA (goat anti-rabbit IgG; Vector Laboratories, USA) then with avidin–biotinylated peroxidase complex diluted 1:500 in PBST (ABC Kit Biotin (ABC; PK-6101, Vector Laboratories, USA). The reaction was visualized using nickel intensified diaminobenzidine (DAB peroxidase substrate kit, Vector Laboratories, USA). Sections from different groups were processed at the same time and using the same conditions and reagents to reduce variability.

### QUANTIFICATION AND STATISTICAL ANALYSIS

#### 3.4 Brain-Wide c-Fos Quantification

38 ± 5.6 coronal sections were selected from each subject for image acquisition, sections covered a large part of the mouse’s brain spacing along the antero-posterior axis from the olfactory bulbs (+2.30 mm relative to bregma) to midbrain (-3.5 mm relative to bregma)^57^. All brain sections were scanned using a Zeiss Axioscan Z1 slide scanner running Zeiss Zen Software (Carl Zeiss MicroImaging, Jena, Germany) with a 20× objective; images were exported in a czi format. Following image acquisition, a small part of the images went through a postproduction refinement using Adobe-Photoshop. Principal Component Analysis (PCA) was applied to compare different staining by using the “dplyr” and the “ggplot2” packages in R (R version 3.4.4, R Development Core Team 2011) which assessed similarity between staining. c-Fos expression and spatial analysis of labelled mice brain section images based on the Allen Brain Atlas were then quantified using the Quint workflow (EBRAINS^37^). The workflow, semi-automated, combines three open-source Software. First, brain section images were correctly aligned to the Allen Mouse Brain Atlas to produce personalized atlas maps fit to match the correct proportions of the sections (QuickNII software). Second, the labelling was segmented from the original images by the Random Forest Algorithm (Ilastik software). Ilastik exports prediction maps which differentiate signal (labelled immunoreactive cells) from background. Finally, the segmented images and customized atlas maps were merged generating quantification data referring to each region present in the reference atlas (Nutil software). The mask feature was also used to define subregions of interest such as the dorsomedial and dorsolateral striatum (DMS, DLS) and the nucleus accumbens core and shell. The mask is applied in addition to, and not instead of, the reference atlas maps.

##### Post processing

Quint provides comprehensive information on all brain regions classified in the Allen Brain Atlas, encompassing a total of 1300 structures. Initially, we excluded data from regions not included in our histochemical procedures—specifically the cerebellum, pons, and olfactory bulbs—resulting in a refined count of 591 regions. Next, we removed regions associated with fiber tracts, bringing the total down to 558 regions. Then we grouped cortical layers as a whole, further reducing the total to 271 regions. Finally, some regions were excluded due to having fewer than two sections in at least six animals, which rendered these data points unreliable. This rigorous selection process ultimately led to a final dataset of 126 regions for analysis.

#### 3.5 Variability

To assess the intra and inter group variability levels and assert statistical relevance a series of analysis were performed. Principal component analysis (PCA) and Heatmaps were first produced. PCA pursues and ranks combinations of variables that account for variance within a given dataset. Both Heatmaps and PCAs were performed using R scripts, the “pheatmap” package for heatmaps and the “dplyr” and “ggplot2” for the PCAs (R version 3.4.4) (R Development Core Team 2011).

##### Standardization

A detailed examination of the z-scores revealed significant variability, particularly within the MC group. This observed variability was likely to introduce substantial bias in the correlations during the network construction phase. To address this issue, we standardized the data by transforming the density of each region for each subject into z-scores. This transformation involved subtracting the mean density across all regions and dividing by the standard deviation, calculated across the same set of regions. The formula used for this standardization is as follows:

Where ***z*** represents the z-score, ***X*** is the raw score (density of each region), *μ* is the mean density across all regions, and *σ* is the standard deviation across all regions. This standardization approach was selected to effectively balance the data, as c-Fos density trends across regions in subjects maintained consistent ratios, thereby preserving the integrity of subsequent correlations.

#### 3.6 Partial Least Square Analysis

Partial Least Squares (PLS) is a powerful multivariate statistical technique utilized in neuroimaging to identify distinct patterns of functional activity or connectivity that differentiate conditions. It leverages singular value decomposition to generate orthogonal latent variable pairs (LV), which capture both contrasts between conditions and brain region saliences. PLS is versatile, applying to both functional activity and regional cell counts, thereby analyzing relationships between brain data and experimental conditions. Statistical significance is determined through permutation testing, while reliability is assessed using bootstrap estimation. Bootstrap resampling with replacement is performed 1000 times to calculate statistical error, allowing saliences to be interpreted as z-values^58,59^. This approach streamlines analysis by avoiding multiple comparison corrections, enabling comprehensive evaluation of all brain regions in a single step. PLS analysis were conducted using a Python script initially developed and provided by the Neuroinformatics Laboratory of Prof. Silvestri at the University of Florence, which we subsequently customized and adapted to meet the specific requirements of our dataset and analysis pipeline.

#### 3.7 Functional connectivity analysis

Within each of the four experimental groups (SS, MS, SC, MC) and Home Cage (HC) animals, all pairwise correlations between c-Fos signal in the 126 regions were determined by computing Pearson correlation coefficients. Each complete set of correlations was computed from a vector of size 8 and were displayed as color-coded correlation matrices using the “psych”, “igraph” and “corrplot” packages in R (R version 3.4.4).

##### Functional network construction

Networks were constructed by thresholding inter-regional correlations in each group. The primary networks were constructed by considering correlations with Pearson’s r > 0.65, which corresponds to a one tailed significance level of P < 0.04, uncorrected for multiple comparisons. Higher and lower confidence networks were constructed as well to ensure that network properties were not dependent on threshold level selection. A threshold of r > 0.71 (corresponding to a significance level of P < 0.025 [one tailed, uncorrected]) was used to generate the high confidence networks and a threshold of r > 0.62 (corresponding to a significance level of P < 0.05 [one tailed, uncorrected]) was used to generate the low confidence networks. In addition, networks were constituted using Spearman’s rank correlation coefficient, with a threshold of r > 0.71, P< 0.025. The nodes in the networks represent brain regions and the correlations that endured thresholding were considered connections. Cytoscape v3.10.0 (http://www.cytoscape.org) was used to visualize networks. In order to evaluate how functional connectivity changed as a function of memory types and acquisition regimen, we categorized our 126 brain regions into 8 taxonomic groups (cortex, olfactory regions, hippocampal formation, cerebral nuclei, thalamus, hypothalamus, midbrain and cortical subplate). While potentially interesting, we did not consider negative correlations in the current network analyses.

##### Scatterplots

We generated scatterplots in *R* (R version 3.4.4, R Development Core Team 2011)) using the “ggplot2” package to assess whether standardized c-Fos density influenced the resulting correlations within our networks.

#### 3.8 Graph theory analysis

Graph theoretical measures were used to characterize properties of each memory and learning regimen networks. For each primary, low and high confidence spatial/cue memory network, 100 random, null hypothesis networks were generated with the same number of active nodes, connections and same degree distribution. Then spatial/cue memory networks’ properties were contrasted with averaged values from these corresponding random, control networks. Network measures and random null hypothesis networks were generated using the “igraph” package in R (R version 3.4.4, R Development Core Team 2011). Definitions and formulae for these graph theory measures are described in the next section. Ninety-five percent confidence intervals for the network measures are reported in order to determine whether network properties differ reliably between experimental groups. Means and confidence intervals for the network measures were derived by bootstrapping which involves resampling subjects with replacement one thousand times and recalculating the network measures.

##### Integration

Integration in a brain network gives rise to coordinated activation of distributed neuronal populations and brain areas. In our analysis we computed two measures of integration: characteristic path length and global efficiency. The shortest path length between two specific nodes is defined as the minimum number of connections that need to be traversed to get from one to the other. The characteristic path length is then the average shortest path length for all pairs of nodes^39^.

The characteristic path length ***L*** can be calculated as:

where:

**N** is the total number of nodes in the network.

**d_ij_** is the shortest path length between nodes **i** and **j.**

**Global efficiency** measures how efficiently information is exchanged over a network. It quantifies the average inverse shortest path length between all pairs of nodes (Rubinov and Sporns, 2010). A higher global efficiency indicates a more efficient network.

##### Segregation

Segregation in a brain network allows for specialized processing in more densely connected clusters. In our analyses we computed two common measures of segregation: mean clustering coefficient and transitivity.

The **clustering coefficient** is computed by dividing the number of existing connections among a node’s directly connected neighbors by the number of possible connections between them. The mean clustering coefficient for the network is then the average clustering coefficient for all the active nodes in the network^39^. The mean clustering coefficient C can be expressed as:

where:

**N** is the total number of nodes in the network.

**C_i_** is the local clustering coefficient for node **i**.

For undirected graphs, the local clustering coefficient for a node v_i_ is given by:

where:

- **T_i_** is the number of triangles connected to node **v_i_**.
- **k_i_** is the degree of node **v_i_**.

**Transitivity** is a weighted version of the clustering coefficient that is less biased by low degree nodes, is defined as the ratio of the number of closed triplets (triangles) to the number of open triplets (connected triples)^60^.

Where **t_i_**is the number of triangles that involve node , and is its degree.

All network metrics, both for integration and segregation and random generated ones, were displayed by using GraphPad Prism 8 (GraphPad Software, Boston, Massachusetts USA).

#### 3.9 Coarse

We categorized nodes into 20 clusters (Table S1). Clusters were delineated accordingly to the Allen Brain Atlas^38^ with few exceptions regarding mainly the frontal cortex, the amygdala, and basal ganglia clusters. Edges between clusters were drawn if there were at least three connections among nodes within clusters. The strength of these edges was determined by the following formula:

Where:

-N: number of valid correlations (r>0.65) between cluster 1 and cluster 2

- n: number of nodes in clusters

-X: cluster 1

-Y cluster 2

For further improved clarity regarding networks interpretation, we have chosen to exclude connectivity arising from connections with the hypothalamic regions at this stage. Networks were visualized using Cytoscape v3.10.0 (http://www.cytoscape.org/).

#### 3.10 Mean R

To evaluate how functional connectivity changed in different memory forms and learning paradigms in our experimental groups, we categorized our 126 brain regions into major brain subdivisions (see Table S1). We then contrasted mean correlation strength between different major subdivisions at both protocol paradigm (SS vs MS, SS-SC vs MS-MC) and memory type (SS-MS vs SC-MC). These contrasts were selected based on coarse grained analysis. Mean correlation strength was determined by the following equation:

- mean corr_ij_: mean correlation between two clusters i and j
- N: total number of possible correlations between clusters i and j.
- Valid corr_ij_: number of correlations between clusters i and j which are r>0.65 and p< 0.04

#### 3.11 Hub

Hub regions play crucial roles in the function of a network. Measures of degree, katz centrality, betweenness centrality and collective influence (CI), were computed for all nodes in networks and used to identify candidate hub regions. **Degree** corresponds to the number of edges a brain region presents in the network.

Node **betweenness** is the number of all shortest paths in the network that pass-through a given node (Brandes, 2001), and nodes with high values of betweenness centrality participate in a large number of shortest paths calculated as:

Where **V** is the set of nodes, σ**(s,t)** is the number of shortest **(s,t)** -paths, and σ**(s,t/v)** is the number of those paths passing through some node **v** other than **s,t**. If **s=t,** σ**(s,t)=1**, and if **v**∈ **s, t,** σ**(s,t/v)=0.**

**Katz centrality** computes the relative influence of a node within a network by measuring the number of immediate neighbors (first degree nodes) as well as all other nodes in the network that connect to the node under consideration through these immediate neighbors (Katz, 1953).

Where **A** is the adjacency matrix of the selected graph (**G**) with eigenvalues λ.

The parameter β controls the initial centrality with the attenuation factor α set at **0.06**, lower than the inverse largest eigenvalue of the adjacency matrix, as required for the Katz centrality to be computed correctly.

Both betweenness centrality and Katz centrality were measured by using “networkX” library in python.

**Collective Influence** is defined as a global centrality measure that calculates the product of the reduced degree of a node and the total reduced degree of all nodes at a specified distance ***d*** from that node^61^. This metric helps identify influential nodes in a network by considering not only the node itself but also its immediate neighbors.

**CI(v)** of a node **v** can be expressed as:

where:

**k(v)** is th degree of node **v**.

- **N_d_(v)** is the set of node that are at distance **d** (set as 4) from node **v.**
- **k(u)** is the degree of node **u**.

CI was produced referring to the “influential” package in R (R version 3.4.4).

- Distance d was set as 4 because it was the measure which most rapidly disrupted network’s GC.

#### 3.11 Network disruption

We assessed network resilience to random removal and targeted attack using an approach previously described by Achard et al (2006)^62^. Random disruption was modeled by sequentially removing nodes and all their connections from the network randomly and then recalculating the size of the largest connected component (i.e., collection of connected nodes) in the new network. Targeted network attacks were simulated using the same process except that node removal began with the most highly ranked node for either node with the highest degree/ Katz centrality/ betweenness centrality/ CI and progressed in order of ranking after recalculating the size of the largest connected component after every removal. Curves describing the effect of random and targeted node removal on relative largest component size were produced for all of the experimental conditions’ memory networks (MC, MS, SS, SC) and compared. Networks’ disruption was determined by applying the “networkX” library in python and represented in GraphPad Prism 8 (GraphPad Software, Boston, Massachusetts USA).

##### Data Collection and Statistical Analysis

Behavioral data were represented as mean ± SEM. One-way or two-way ANOVA as well as post hoc analysis were performed using PRISM 10 software (were performed using PRISM 10.6 software (GraphPad Software, Boston, Massachusetts USA).

