## Supplemental Text and Figures for "Time between learning events shapes functional connectivity in spatial memory, a brain-wide analysis"

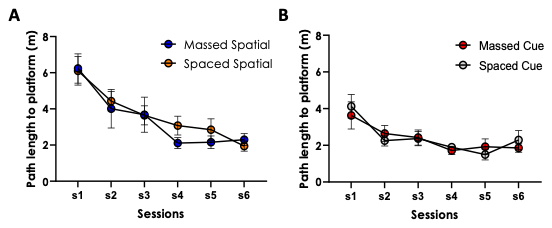
**Suppl. Figure 1**. **Path Length Reduction Across Training Sessions in Massed and Spaced Groups.** Path length to reach the platform across training sessions (mean +/- S.E.M.). **(A)** Both massed and spaced groups in the spatial version of the MWM showed progressive decreases in the path length to find the platform, with no significant differences between protocols (two-way repeated measures ANOVA: session F(5,110) = 11.58, p < 0.0001; protocol F(1,22) = 0.3252, p = 0.5743; session × protocol F(5,110) = 0.2093, p = 0.9579). The one-way ANOVA on distance traveled to reach the platform confirmed that both groups successfully learned the task path length: massed spatial F(5,54) = 5.068, p = 0.0007; spaced spatial F(5,72) = 6.660, p < 0.0001). **(B)** Decreased traveled distance was observed across sessions in both massed and spaced cue groups, with no significant differences between protocols (two-way repeated measures ANOVA: session F(5,115) = 7.710, p < 0.0001; protocol F(1,23) = 0.01686, p = 0.8978; session × protocol F(5,115) = 0.4878, p = 0.7848). The one-way ANOVA on distance traveled to reach the platform confirmed that both groups successfully learned the task path length: massed cue F(5, 66) = 2.583, p = 0.0340; spaced cue F(5,72) = 4.717, p = 0.0009).

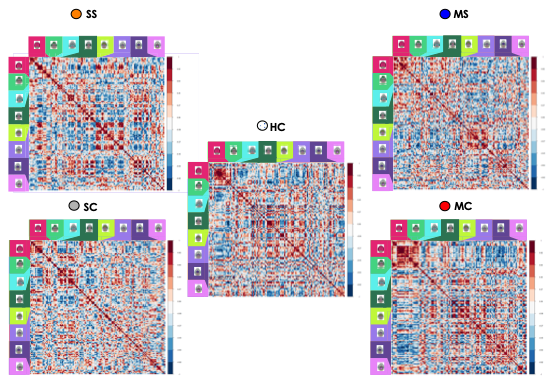

**Suppl. Figure 2. Functional connectivity matrices based on c-Fos co-activation.** Matrices present pairwise Pearson correlation coefficients among the 126 brain regions for each experimental group (SS, MS, SC, MC) and home cage controls (HC). Colors reflect correlation strength (scale on the right) ranging from dark blue (strong negative correlations) to dark red (strong positive correlations). Brain regions were divided into 8 major taxonomy groups: cortex (magenta), olfactory regions (light green), hippocampal formation (cyan), cortical subplate (dark green), cerebral nuclei (lime), thalamus (lavender), hypothalamus (purple), and midbrain (violet). The matrices reveal network organization differences among spaced and massed protocols, as well as between spatial and cue learning tasks. Spaced Spatial (SS); Massed Spatial (MS); Spaced Cue (SC); Massed Cue (MC); Home Cage controls (HC).

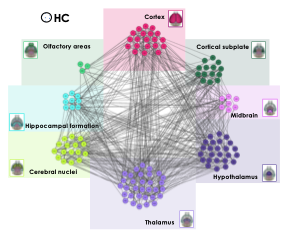

**Suppl. Figure 3. Home Cage (HC) functional connectivity network based on c-Fos co-activation.** Network graph generated from significant positive inter-regional correlations (Pearson’s r > 0.65, p < 0.04) for the HC group. The 126 brain regions are divided into 8 taxonomic groups: cortex (magenta), olfactory regions (light green), hippocampal formation (cyan), cortical subplate (dark green), cerebral nuclei (lime), thalamus (lavander), hypothalamus (violet), and midbrain (pink).

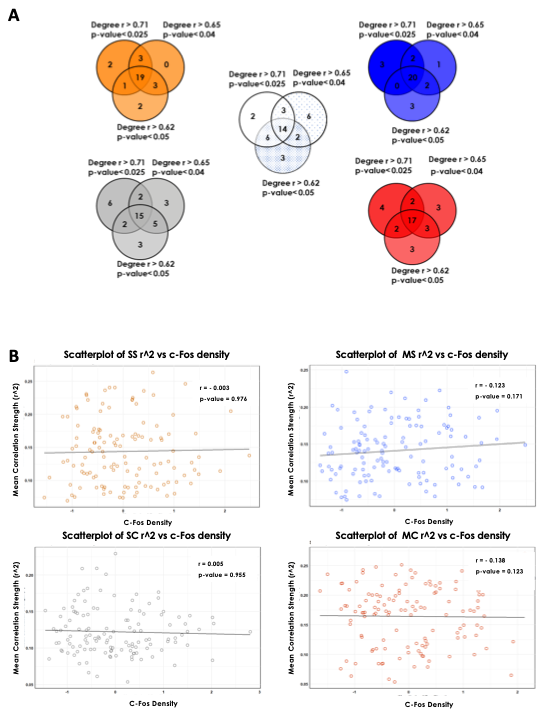

**Suppl. Figure 4. Network Node Degree Stability and c-Fos Density Influence on Functional Connectivity. A.** Venn diagram representing correlation-based network node degree comparison derived from Pearson coefficients at different thresholds (r > 0.62, 0.65, 0.71), showing consistent degree patterns across thresholds for all experimental and HC groups. **B.** Scatterplots assessing the relationship between c-Fos density and squared correlation strength (r²) for each experimental group, reporting the following statistics: (SS): r = -0.003, p = 0.976; (MS): r = -0.123, p = 0.171; (SC): r = 0.005, p = 0.955; (MC): r = -0.138, p = 0.123. These non-significant correlations indicate that regional c-Fos density does not bias correlation strength in the constructed functional networks.

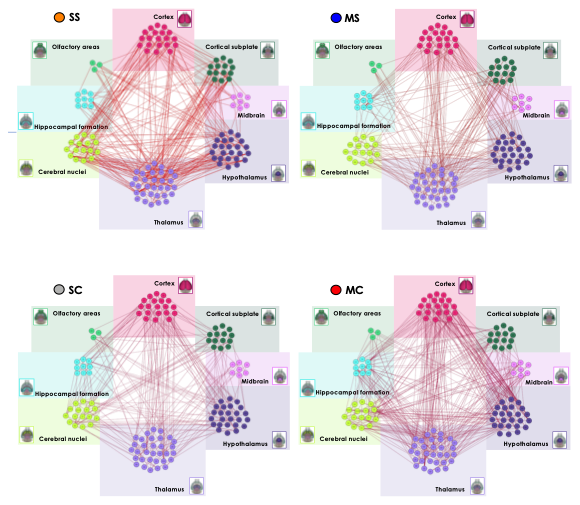

**Suppl. Figure 5. Functional negative connectivity networks based on c-Fos co-activation.** Network graphs generated by considering only significant negative inter-regional correlations (Pearson’s r < -0.65, p < 0.04) for the spaced spatial (SS), the massed spatial (MS), the spaced cue (SC) and massed cue (MC) groups. The 126 brain regions are clustered in 8 taxonomic groups: cortex (magenta), olfactory regions (light green), hippocampal formation (cyan), cortical subplate (dark green), cerebral nuclei (lime), thalamus (lavander), hypothalamus (violet), and midbrain (pink). Spaced Spatial (SS); Massed Spatial (MS); Spaced Cue (SC); Massed Cue (MC).

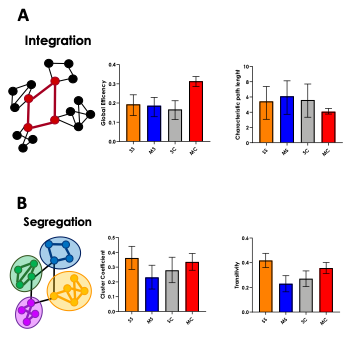

**Suppl. Figure 6. Key global network metrics of integration and segregation across experimental groups.** Bar graphs show integration and segregation metrics across experimental groups (SS, MS, SC, MC). **A.** Integration Measures: Global efficiency and characteristic path length reflect the efficiency and capacity for small world network communication. **B.** Segregation Measures: Clustering coefficient and transitivity indicate the extent of local clustering, community structure, and preferential connectivity between nodes of similar degree. All groups exhibit similar small-world network attributes, with no significant differences emerging between spaced and massed protocols or task types. Error bars represent 5-95% confidence intervals evaluated upon 1000 iterations. Spaced Spatial (SS); Massed Spatial (MS); Spaced Cue (SC); Massed Cue (MC).

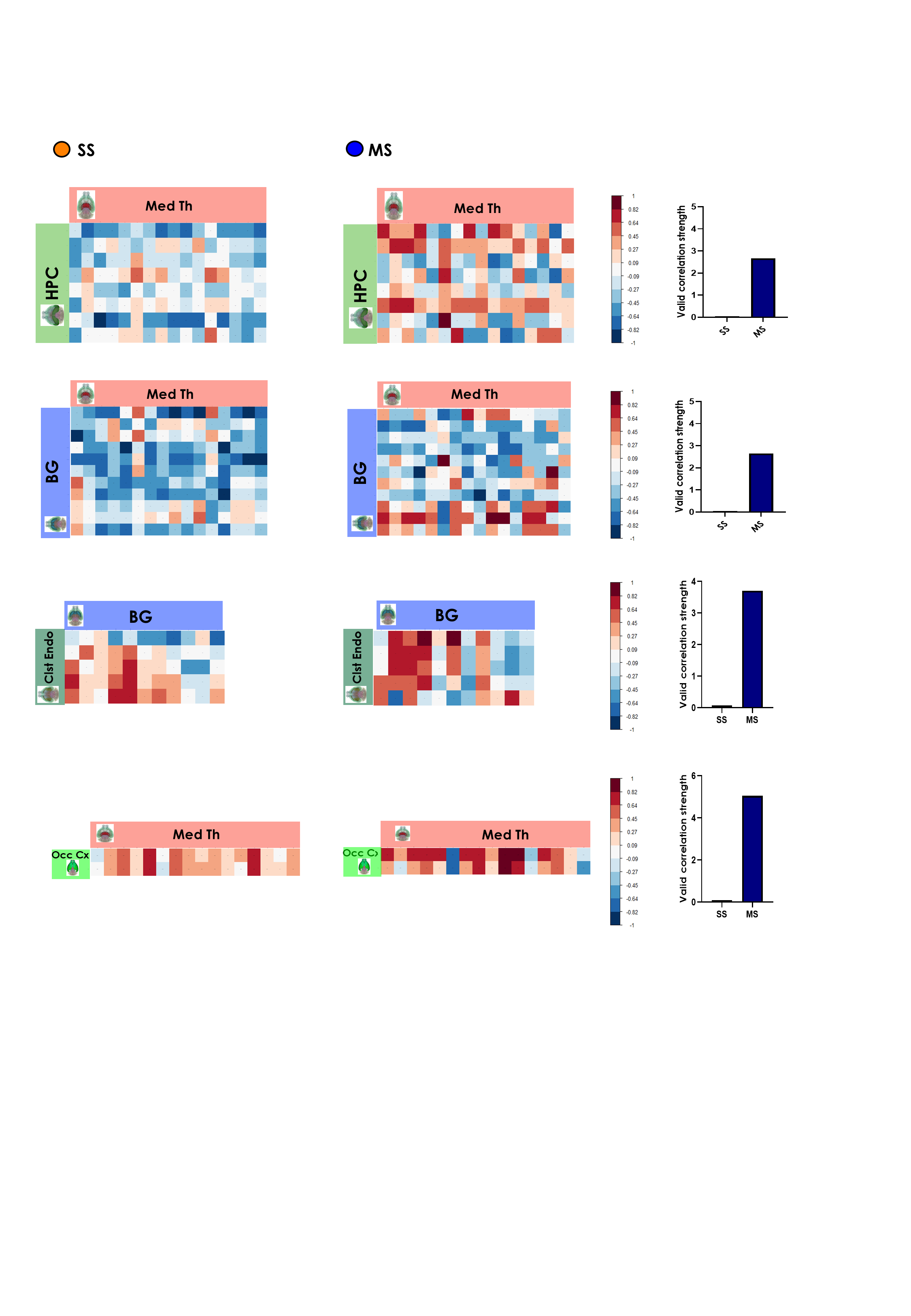

**Suppl. Figure 7. Functional connectivity differs as a function of the training protocol: clusters showing higher correlation in MS compared to SS**. Color-coded matrices showing inter-regional correlations for Fos expression after spaced and massed spatial training. Histograms represent correlation strength between clusters. HPC (hippocampus), Med Th (medial thalamus), BG (basal ganglia), Clst Endo (claustrum-endopiriform cortex), Occ Cx (occipital cortex). Spaced Spatial (SS); Massed Spatial (MS).

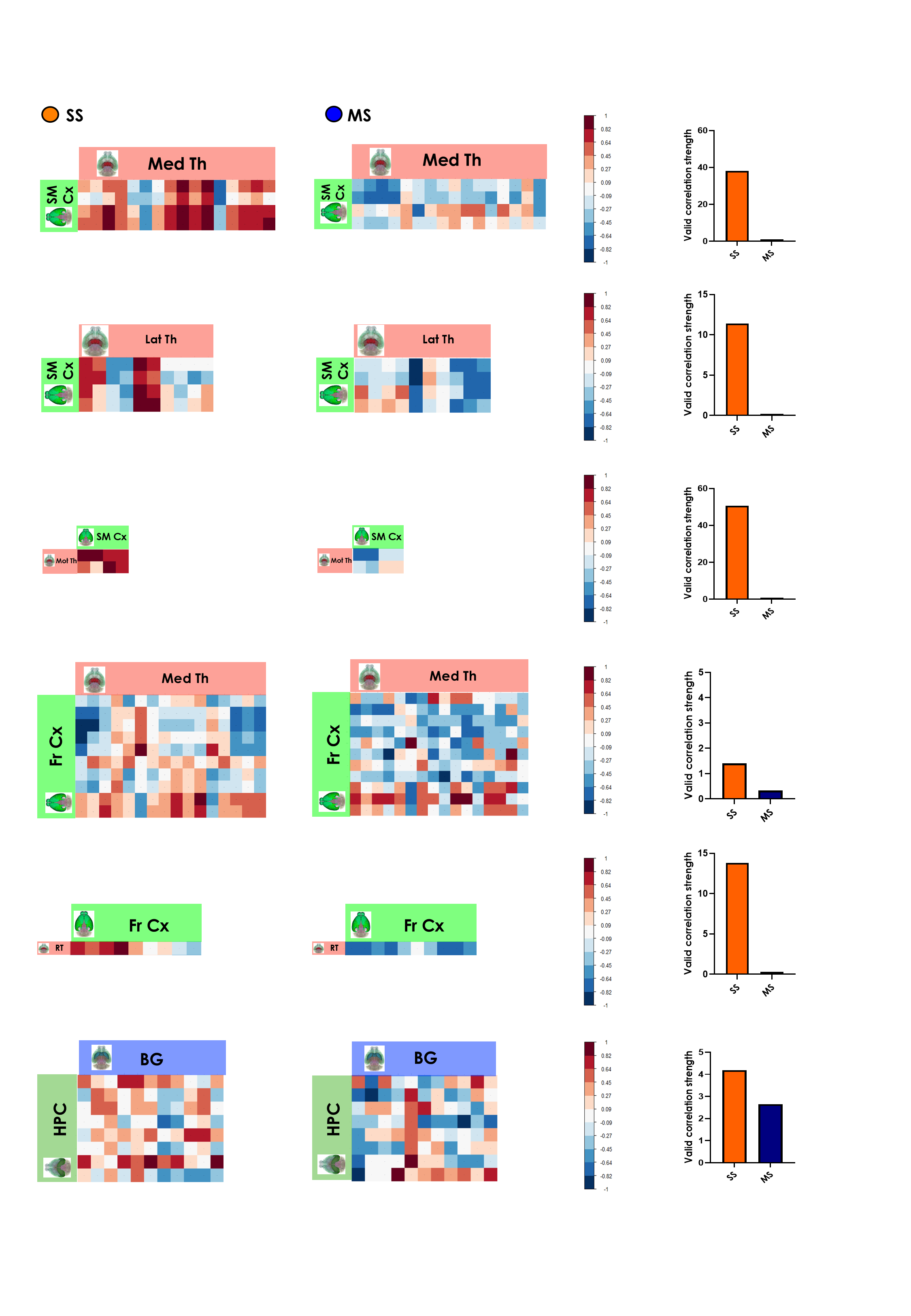

**Suppl. Figure 8.** **Functional connectivity changes as a function of the training protocol: clusters showing higher correlation in SS compared to MS**. Color-coded matrices showing inter-regional correlations for Fos expression after spaced and massed spatial training. Histograms represent correlation strength between clusters. SM Cx (sensory-motor cortex), Med Th (medial thalamus), Lat Th (lateral thalamus), Mot Thal (motor thalmus), Fr Cx (frontal cortex), RT (reticular nucleus of the thalamus), BG (basal ganglia), HPC (hippocampal complex). Spaced Spatial (SS); Massed Spatial (MS).

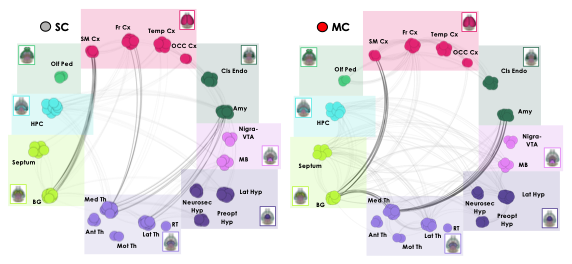

**Suppl. Figure 9. Coarse grained functional connectivity networks in spaced cue (SC) and massed cue (MC) training groups.** Brain networks, based on c-Fos correlation patterns. The 126 brain regions divided into 8 taxonomic groups, were organized into 20 functional-clusters (cortex (magenta): somato-motor cortex (SM Cx), frontal cortex (Fr Cx), temporal cortex (Temp Cx) and occipital cortex (OCC Cx). Olfactory regions (light green): olfactory peduncle (Olf Ped). Hippocampal formation (cyan): hippocampus (HPC). Cerebral nuclei (lime): Septum and basal ganglia (BG). Thalamus (lavender): medial thalamus (Med Th), anterior thalamus (Ant Th), motor thalamus (Mot Th), lateral thalamus (Lat Th), reticular thalamus (RT). Hypothalamus (purple): neurosecretory hypothalamus (Neurosec Hyp), preoptic hypothalamus (Preopt Hyp), and lateral hypothalamus (Lat Hyp). Midbrain (violet): nigra and ventral tegmental area (Nigra-VTA) and midbrain regions (MB). Cortical subplate (dark green): claustrum and endopiriform regions (Cls Endo) and amygdala (Amy). Edges are represented as connections between clusters only if there were at least three significant inter-cluster connections, highlighting robust communication. Darker lines denote interesting relevant correlations. **A.** SC exhibited strong functional connectivity between the somatomotor cortex (SM Cx) and basal ganglia, as well as between the frontal cortex and medial thalamus. Additionally, strong connectivity was observed between the amygdala and both the medial and lateral thalamus (med th and lat th). **B.** MC showed strong correlations between the medial thalamus and the amygdala, as well as between the medial thalamus and the basal ganglia (BG), which correlate with the somatomotor cortex (SM Cx**).**

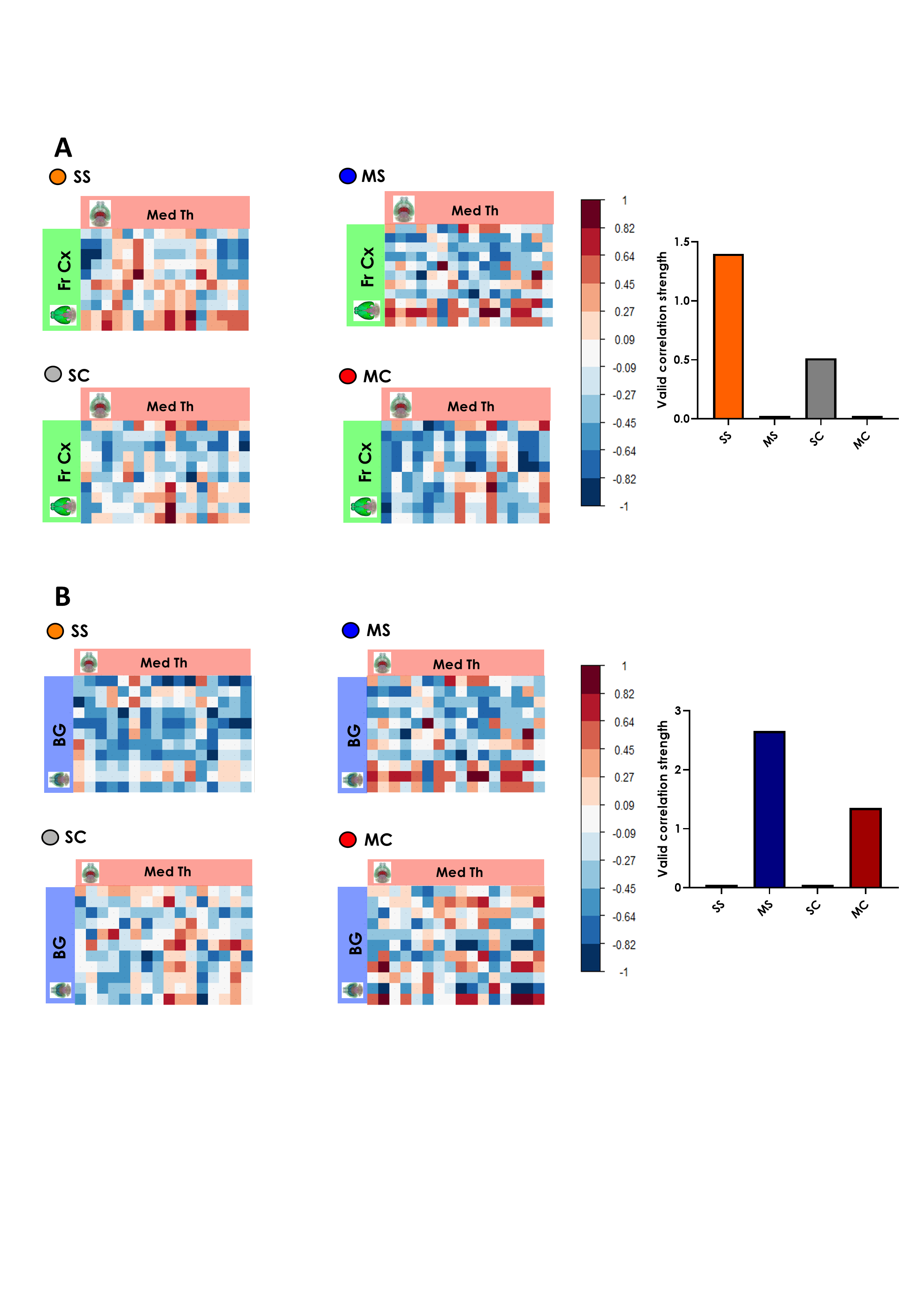

**Suppl. Figure 10. Functional connectivity changes as a function of the training protocol**. Color-coded matrices showing inter-regional correlations for Fos expression between in the 4 experimental groups: Spaced Spatial (SS); Massed Spatial (MS); Spaced Cue (SC); Massed Cue (MC). **A.** Frontal cortex and the medial thalamic clusters show a higher correlation after spaced compared to massed training independently on the training protocol. **B.** The basal ganglia and the medial thalamic clusters show higher correlation in the massed compared to the spaced trained group independently of the training protocol. Histograms represent correlation strength between clusters. Fr Cx (frontal cortex cluster), Med Th (medial thalamus cluster), BG (basal ganglia cluster).

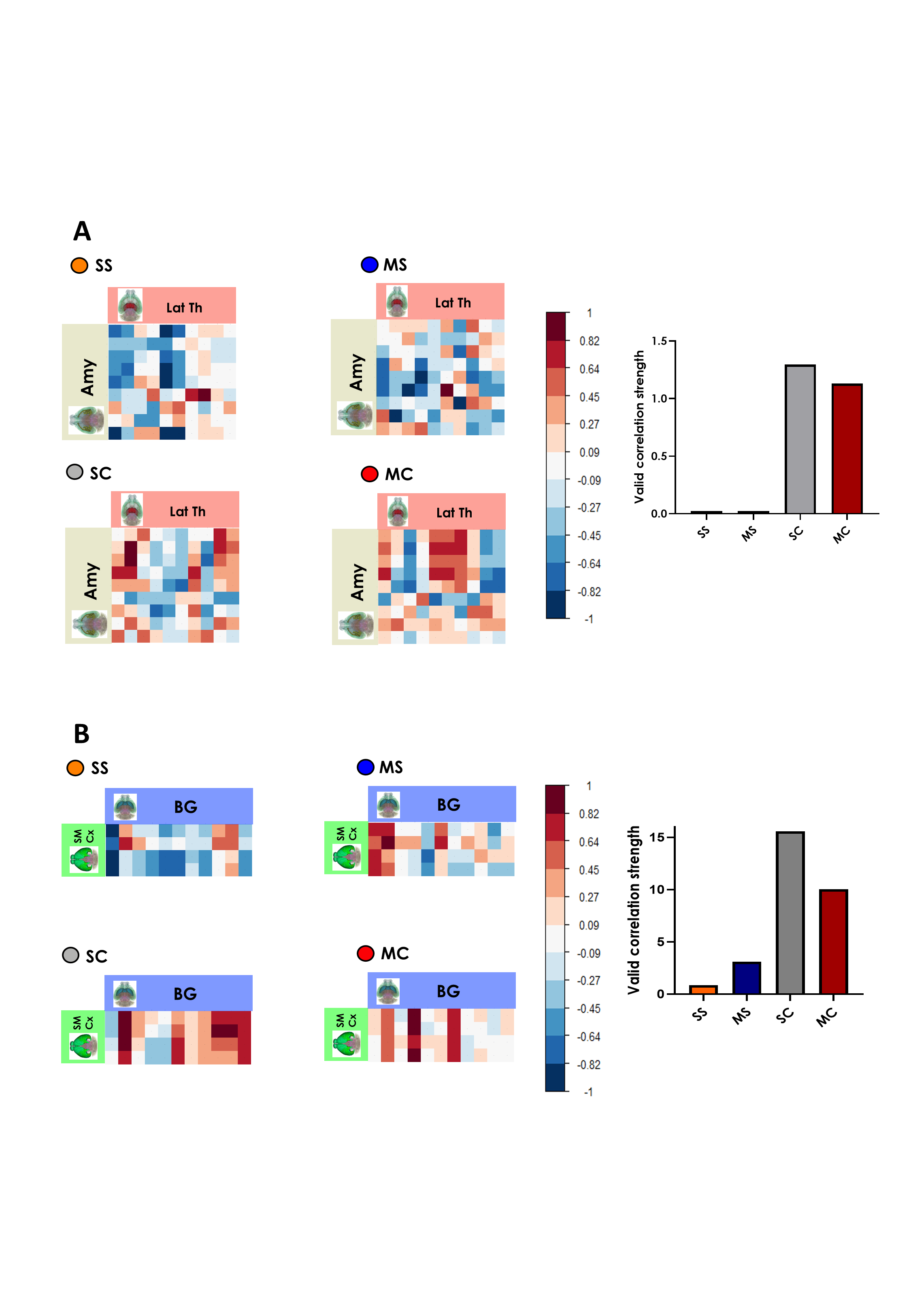

**Suppl. Figure 11. Functional connectivity changes as a function of the type of memory: clusters showing higher correlation in the cue compare to spatial trained groups**. Color-coded matrices showing inter-regional correlations for Fos expression between in the 4 experimental groups: Spaced Spatial (SS); Massed Spatial (MS); Spaced Cue (SC); Massed Cue (MC). **A.** The amygdala and the lateral thalamic clusters show a higher correlation in the cue compared than in the spatial training group independently on the training protocol. **B.** The basal ganglia and the sensory motor cortical clusters show higher correlation in the cue compared than in the spatial training group independently on the training protocol. Histograms represent correlation strength between clusters. Lat Th (lateral thalamus cluster), Amy (amygdala cluster) SM Cx (sensory motor cortex cluster), BG (basal ganglia cluster).

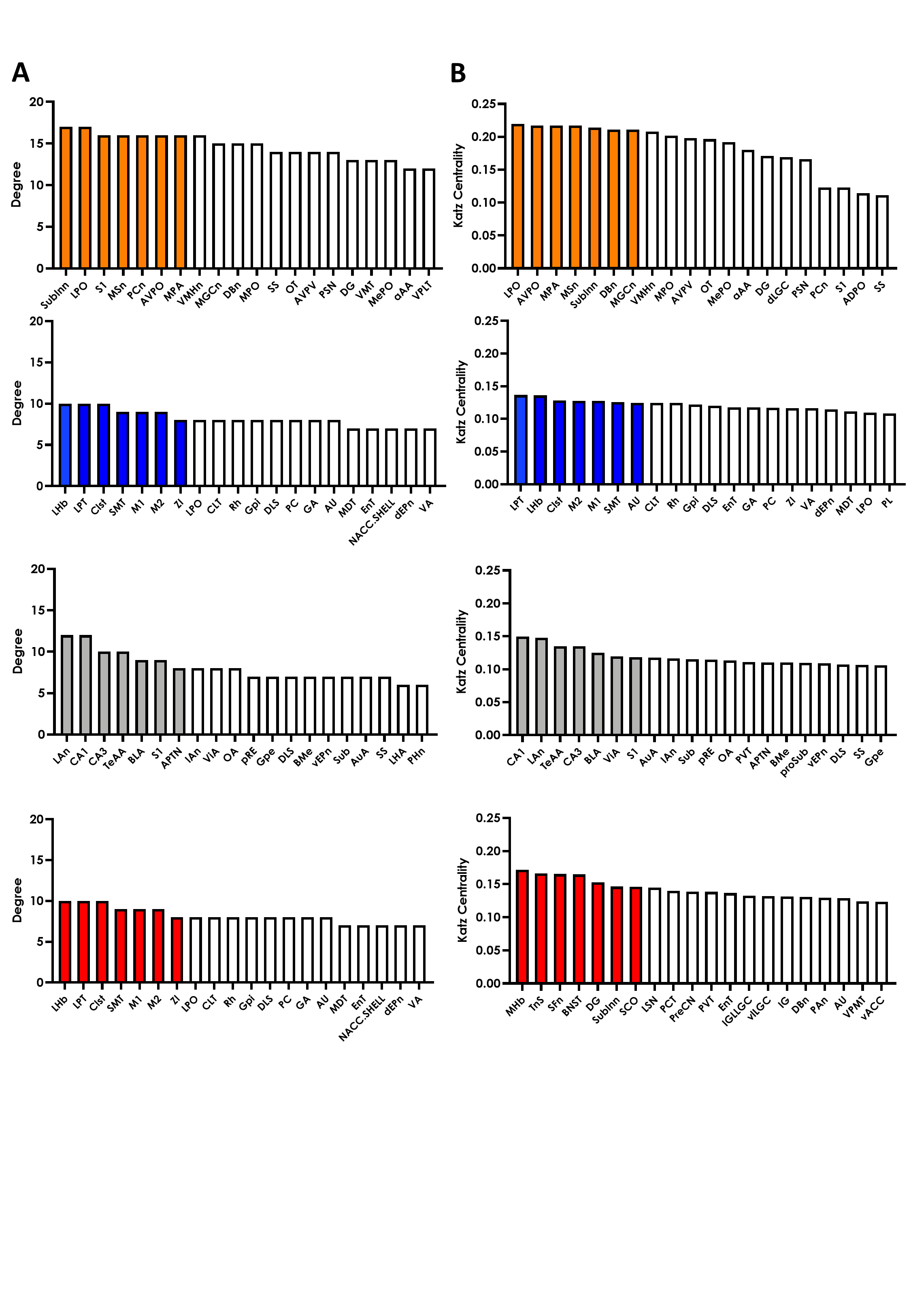

**Suppl. Figure 12. Ranking of the most disruptive nodes in all experimental groups based on degree and Katz centrality.** Disruption ranking of nodes in the spaced spatial (SS, orange), massed spatial (MS, blue), spaced cue (SC, grey) and massed cue (MC, red) networks derived from simulated node removal using **A.** degree and **B.** Katz centrality. The top 7 nodes for each group are highlighted in their respective colors; all other nodes are shown in white.

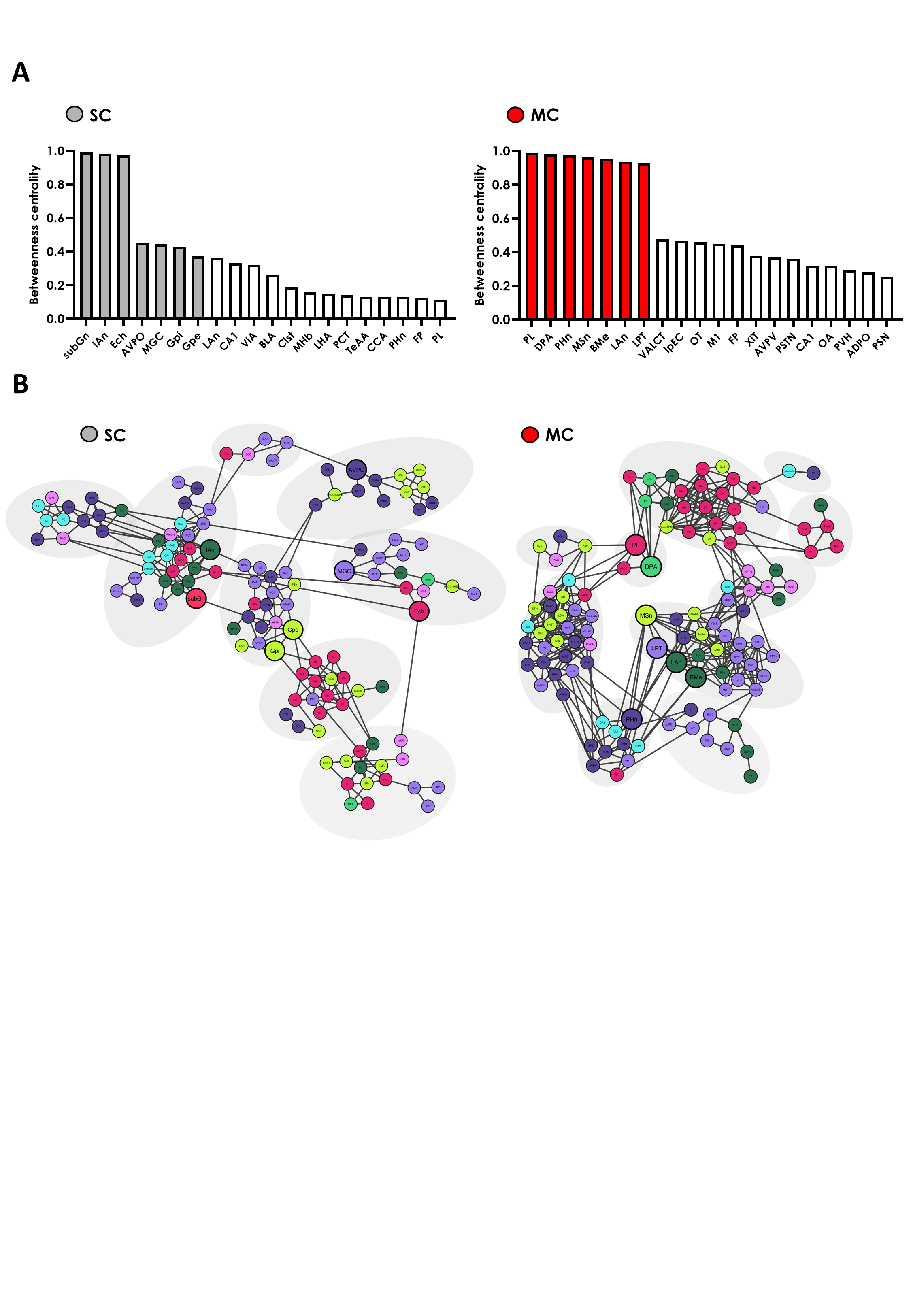

**Suppl. Figure 13. Ranking and network visualization of the most disruptive nodes in spaced and massed cue networks based on betweenness centrality. A.** Disruption ranking of nodes in the spaced cue (SC, grey) and massed cue (MC, red) networks derived from simulated node removal using betweenness centrality. The top 7 nodes for each group are highlighted in their respective colors; all other nodes are shown in white. **B.** Network layout visualizations generated with Cytoscape for SC and MC groups. Grey ellipses highlight networks’ communities. Nodes are color-coded according to their original taxonomic groups: cortex (magenta), olfactory regions (light green), hippocampal formation (cyan), cortical subplate (dark green), cerebral nuclei (lime), thalamus (lavander), hypothalamus (purple), and midbrain (violet). The top 7 ranked disruptive nodes (from panel A) emphasized by larger node sizes to highlight their network importance.
